# Paralemmin-3 sustains the integrity of the lateral plasma membrane and subsurface cisternae of auditory hair cells

**DOI:** 10.1101/2025.07.11.664127

**Authors:** Victoria C. Halim, Iman Bahader, Dennis Derstroff, Christina Ullrich, Makoto F. Kuwabara, Greta Hultqvist, Kathrin Kusch, Loujin Slitin, Lore Becker, Martin Hrabè de Angelis, Carolin Wichmann, Dominik Oliver, Nicola Strenzke, Christian Vogl, Manfred W. Kilimann

**Author notes:** Correspondence should be addressed to: Christian Vogl or Manfred W. Kilimann. these authors contributed equally (Christian Vogl, Manfred W. Kilimann).

## Abstract

In the mammalian inner ear, cochlear inner hair cells (IHCs) enable accurate and faithful synaptic sound encoding, while outer hair cells (OHCs) perform frequency-specific sound amplification and fine-tuning through their intrinsic voltage-dependent somatic electromotility. This latter process is facilitated by the unique trilaminate structure of the OHC lateral wall, which consists of the plasma membrane that is densely occupied by the transmembrane motor protein Prestin, the submembrane actin- and spectrin-based cytoskeleton, and the endomembranous subsurface cisternae. This complex system provides mechanical resilience while allowing for cell expansion and contraction during electromotility. Whereas the ultrastructure of the lateral wall is well described, its molecular architecture remains largely elusive. Here, we identified Paralemmin-3 (Palm3) as a novel protein specifically localized to the lateral walls of auditory HCs to play a crucial role in connecting the plasma membrane to the underlying cytoskeleton and subsurface cisternae. *Palm3*-KO mice display early-onset and progressive hearing impairment that results from diminished cochlear amplification. Subsequent multiscale morphological analyses revealed structural collapse of OHCs that led to progressive and extensive OHC loss along the tonotopic axis. *Palm3*-KO OHCs exhibited disrupted expression and distribution of several membrane-associated proteins – including spectrin isoforms and Prestin – suggesting an essential role of Palm3 in plasma membrane scaffolding. Electron tomography of OHC lateral walls revealed significantly fewer and structurally perturbed subsurface cisternae. Finally, adeno-associated virus (AAV)-mediated rescue of Palm3 during early postnatal development partly restored hearing function, enhanced OHC survival, and restored OHC cell shape as well as membrane protein expression levels. In summary, Palm3 is a key component of the submembrane cytoskeleton in cochlear hair cells, playing a fundamental role in hair cell biology and hearing, and emerges as an attractive candidate for the long-elusive “pillar” component of the hair cell lateral wall ultrastructure.

## Introduction

In the mammalian inner ear, two functionally distinct populations of auditory hair cells translate physical sound waves into neural code: whereas inner hair cells (IHCs) faithfully encode sound at their specialized ribbon synapses with afferent spiral ganglion neurons, outer hair cells (OHCs) enable the exquisite frequency discrimination capacity of the mammalian cochlea by actively amplifying basilar membrane motion through their intrinsic voltage-dependent somatic electromotility. The capacity of OHCs to undergo rapid, sub-millisecond length changes requires a delicate balance between radial stiffness and axial compliance to sustain repeated cycles of elongation and contraction without loss of structural integrity during sound transduction (Borkó *et al*, 2005; Batta *et al*, 2003; Hallworth, 1995; He & Dallos, 2000). This equilibrium between flexibility and rigidity is believed to arise from the unique trilaminate architecture of the OHC lateral membrane, which serves as the foundation for the mechanical resilience of OHCs.

The OHC plasma membrane (PM) is densely decorated with intramembrane particles – subsequently identified as Prestin motor proteins – that mediate voltage-dependent conformational changes to drive somatic electromotility (Gulley & Reese, 1977; Kalinec *et al*, 1992; Zheng *et al*, 2000). Beneath the PM lies the cortical lattice, a meshwork of actin filaments and spectrin cross-links that maintains OHC shape and modulates length and stiffness in response to mechanical forces (Holley *et al*, 1992; Holley & Ashmore, 1988, 1990; Holley, 1996; Legendre *et al*, 2008). Layered underneath the lattice, the subsurface cisternae (SSC) form a circumferential lamellar system (Forge *et al*, 1993), whose cell-biological function is largely unclear but likely includes (i) the maintenance of potential gradients (Spoendlin, 1966), (ii) submembraneous Ca^2+^ homeostasis (Ikeda & Takasaka, 1993; Fuchs *et al*, 2014) and (iii) optimization of motor protein activation (Song & Santos-Sacchi, 2015). Connecting the plasma membrane with the OHC submembrane cytoskeleton are electron-dense structures previously referred to as ‘pillars’ (Bannister *et al*, 1988; Flock *et al*, 1986; Forge, 1991; Holley *et al*, 1992; Holley & Ashmore, 1988). On ultrastructural level, pillars are regularly spaced (26 ± 11 nm) but vary in length (17-51 nm), orientation, and often bifurcate into a ‘Y’-shape at their inner end, with a diameter of 3-4 nm (Forge, 1991; Holley *et al*, 1992; Triffo *et al*, 2019). This structural variability suggests dynamic conformations, consistent with a role as entropic springs within the OHC trilaminate structure (Raphael, 2022). Although first described by electron microscopy in the late 1980s, the molecular identity of the pillars has remained elusive.

Paralemmins are a protein family of four isoforms: Paralemmin-1 to Paralemmin-3 (Palm1-3) and Palmdelphin (Palmd). In the course of evolution, the paralemmin family appeared with the emergence of vertebrates (Hultqvist *et al*, 2012). The isoforms share a molecular layout consisting of an N-terminal coiled-coil domain and a C-terminal CaaX motif which is lipidated to constitute a membrane anchor. These highly conserved features are connected by intervening sequences which are more divergent between the four isoforms and are predicted to be largely intrinsically unstructured. Early observations suggested an involvement of paralemmins in plasma membrane dynamics and cell shape control (Kutzleb *et al*, 1998). More recently, interactions with the actin cytoskeleton have emerged for Palmd (Kalebic *et al*, 2019; Sáinz-Jaspeado *et al*, 2021), and with spectrin for Palm1 (Macarrón-Palacios *et al*, 2025). Palm1, in particular, was identified as a powerful regulator of the actin/spectrin-based membrane-associated periodic skeleton (MPS) of neuronal axons. The biological roles and molecular properties of Palm2 and Palm3 remain poorly understood.

*Palm3*-deficient mice were generated in the context of the EUCOMM mutagenesis programme and were subjected to the phenotyping screen of the German Mouse Clinic (GMC). While homozygous *Palm3*-mutant mice were apparently healthy and fertile, and had normal outcomes in most phenotyping parameters, they displayed reduced startle reflex and pre-pulse inhibition in behavioral testing, and impaired clickbox and auditory brainstem response (ABR) in neurological analysis. These observations suggested hearing impairment, and the gEAR single-cell RNAseq database (Orvis *et al*, 2021) indicated abundant expression of Palm3 mRNA in both populations of cochlear hair cells but in no other cell population of the organ of Corti. Given the marked cell-biological demand of auditory hair cells for durable yet flexible membrane scaffolding, especially in electromotile OHCs, and the recently identified role of Palm1 in organizing the submembrane cytoskeleton and anchoring it to the plasma membrane, we hypothesized that Palm3 might be crucial for the mechanical resilience of hair cells by mediating, or contribution to, the membrane-anchoring of their submembrane skeleton.

Therefore, we set out to characterize Palm3 distribution in the murine cochlea and investigated the morphological and functional consequences of *Palm3* deficiency in the peripheral auditory system using systems-physiology, immunohistochemistry, and advanced microscopy techniques – including lightsheet, confocal, 2D-STED nanoscopy, and electron tomography. *Palm3*-KO mice exhibited early-onset, progressive hearing loss due to severe degeneration of auditory hair cells across the entire tonotopic axis. Compatible with a role in membrane scaffolding, Palm3 specifically concentrates in the lateral membranes of both IHCs and OHCs, where it colocalizes with spectrins. *Palm3* loss from OHCs resulted in drastically shortened cell length, reduced spectrins membrane-expression, and compromised subsurface cisternae integrity. Notably, AAV-mediated delivery of wildtype *Palm3* into *Palm3*-KO mice partially restored auditory function, OHC morphology and spectrin localization.

In summary, our data identifies *Palm3* as a novel putative deafness gene and Palm3 as a promising protein candidate contributing to the proposed ‘pillar complexes’ that link the submembraneous cytoskeleton to the plasma membrane in auditory hair cells.

## Materials and Methods

### Animals

All animal experiments were conducted according to national, regional and institutional guidelines of either the University Medical Center Göttingen, Lower Saxony, Germany or the Medical University Innsbruck, Tyrol, Austria. *Palm3*^-/-^ mice were obtained through either homozygous (*Palm3*^-/-^) or heterozygous (*Palm3*^+/-^) breeding pairs. Wild-type control (*Palm3*^+/+^) mice were obtained from heterozygous (*Palm3*^+/-^) or C57BL/6N breeding pairs. All animals were kept in individually ventilated cages within a specific pathogen-free facility on a 12-h light/dark cycle. Animals were housed in social groups and had unrestricted access to food and water. Male and female mice were included in the study, and no significant sex-based differences were observed.

### Generation of *Palm3*-KO mice

Palm3-mutant mice were generated in the EUCOMM mutagenesis programme by targeted insertion of a *lacZ* reporter gene, a *neo* selection gene, and an array of *frt* and *loxP* cassettes into introns 1 and 2 of the Palm3 gene of C57BL/6N ES cells. Blastocyst injection of ES cell clone EPD0230_4_A01 at the European Mouse Mutant Archive (EMMA; Monterotondo, Italy) generated the conditional mouse line *Palm3*^tm1a(EUCOMM)Wtsi^. Allele design and availability of mice and materials can be found on the web-page of the International Mouse Phenotyping Consortium (IMPC): https://www.mousephenotype.org/data/alleles/MGI:1921587/tm1a(EUCOMM)Wtsi. Frozen sperm from these mice were imported from EMMA Monterotondo into the Transgenic Facility of the Max-Planck-Institute for Multidisciplinary Sciences (City Campus) and employed for in-vitro fertilization of eggs from transgenic mice expressing Cre recombinase under the control of the EIIa promoter. Deletion of exon 2 together with the *neo* transgene between the two outer-most *loxP* cassettes in founder progeny, generating the constitutive KO allele *Palm3*^tm1b^, was confirmed by PCR and sequencing. For genotyping, the primer pair 5’-CTCTAAGTCCACAACTAGGCAGGA-3’ (Sense_Upstream-Exon2-38212) and 5’-GCTTGACCCCAATTGTGC-3’ (Asense_Upstream-Exon2-38213) detects the WT allele (340 bp amplimer), whereas the primer pair 5’-CACTGCATTCTAGTTGTGGTTTGT-3’ (Sense_SV40pA_03822) and 5’-GAGCTTGGGTTGCACTGGTC-3’ (Asense_Downstream-Exon2-38214) detects a 146 bp KO amplimer after deletion of the *neo*/exon2 sequence block. Exon 2 deletion causes a reading-frame shift but retains the *lacZ* reporter cassette. The EIIa-*Cre* transgene was subsequently bred out, and the colony was maintained in the C57BL/6N background.

### Auditory systems physiology

ABR and DPOAE measurements were performed as previously described (Jing *et al*, 2013; Strenzke *et al*, 2016; Pauli-Magnus *et al*, 2007). ABR and DPOAE measurements for the initial phenotype description of *Palm3*-KO were performed in 3-week- and 12-13-week-old *Palm3*-KO and WT littermates, whereas measurements for *Palm3*-rescue experiments were performed in AAV-injected 5–6-week-old *Palm3*-KO mice. Animals were anesthetized by intraperitoneal injection of ketamine (125mg/kg) and xylazine (2.5 mg/kg). The heart rate was continuously monitored throughout the procedure. A rectal temperature-controlled heat blanket (Hugo Sachs Elektronik–Harvard Apparatus) was used to maintain a constant core temperature of 37℃. Stimulus generation, presentation, and data acquisition was carried out using the TDT Systems III operated by BioSig32 software (Tucker Davis Technologies). Acoustic stimulus in form of either 12 ms tone-burst (10 ms plateau, 1 ms cos2 rise/fall) or clicks of 0.03 ms were presented ipsilaterally at a stimulation rate of 40 Hz and 20 Hz, respectively, at frequencies of 4, 6, 8, 12, 16, 24, and 32 kHz in the free field using a JBL 2402 speaker (JBL GmbH & co, Neuhofen, Germany). ABRs were recorded by placing subcutaneous needle electrodes at the vertex and near the mastoid region, with a ground electrode positioned dorsally near the base of the tail. The difference potential between the recording electrodes was amplified (50,000x using a custom-built JHM NeuroAmp 401 amplifier), filtered (0.4-4 kHz), and averaged over 1300 sweeps to generate two mean traces per sound intensity. Recording contaminated by noise, primarily due to ECG interference, were automatically excluded using the artifact rejection function in BioSig32. Click ABRs were measured in 10 dB SPL steps. ABR threshold was subjectively determined as the lowest stimulus intensity eliciting a consistent waveform in both averaged traces and were validated by an independent observer. ABR peaks were manually assigned according to their normal latency values, and each wave amplitude was calculated as the difference between highest point of the wave and the subsequent local minimum. In rescue experiments, recordings were first performed on the AAV-injected (ipsilateral) ear of *Palm3*-KO mice, then occluded using electrode gel and tissue to induce a 30-40 db conductive hearing loss (Pauli-Magnus *et al*, 2007) to avoid acoustic crosstalk for the subsequent measurements in the non-injected (contralateral) ear. Tone burst thresholds that surpassed 80 dB were assigned as 90 dB for the purpose of calculating means and SEMs. For DPOAE measurements, two continuous sine wave stimuli (f1, f2) were generated using Tucker-Davis system III and ED1/EC1 speaker system (Tucker-Davis). A custom-built ear probe was employed to deliver two primary tones into the ear canal and to record DPOAEs using a miniature microphone (MKE-2, Sennheiser, Hannover, Germany). The microphone signal was amplified via an external sound card (Terratec DMX 6Fire USB) and digitized with a TDT III system. All systems-physiological experiments were performed in compliance with the national animal care guidelines and were approved by both the board for animal welfare of the University Medical Center Göttingen and the State Animal Welfare office of Lower Saxony (AZ19/3133).

### Immunohistochemistry

Cochleae were fixed with 4% FA in PBS for 1h on ice. Apical turns of the organ of Corti were dissected out of the cochleae and treated with 0.5% Triton-X100 in PBS for permeabilization. This was followed by a blocking step using a PBS-based blocking buffer (BB) containing 0.5% Triton-X100 and 10% goat serum for 1h at RT. Subsequently, samples were incubated overnight with primary antibodies diluted in BB. The following primary antibodies were used: mouse anti-alpha2 Spectrin (SH3-domain, #803201, Biolegend), mouse anti-beta1 Spectrin (aa 1555-2137, #75-374, Antibodies inc), mouse anti-beta2 Spectrin (aa 2101-2189, #612562, BD Transduction Laboratories™), chicken anti-Calretinin (#214 106, Synaptic Systems), mouse anti-HuD (H-9) (#sc-48421, Santa Cruz), mouse anti-Myosin7a (#MYO7a 138-1, Developmental Studies Hybridoma Bank), guinea pig anti-Parvalbumin (#195 004, Synaptic Systems), and rabbit anti-Prestin (#ab242128, Abcam). The next day, secondary antibodies and/or nanobodies were added to the samples. The following secondary antibodies were used: anti-GFP FluoTagX4 Atto488 (#N0304-At488-L, NanoTag), anti-mouse FluoTagX2 IgG1 488 (#N2002-At488, NanoTag), anti-mouse FluoTagX2 IgG1 635p (#N2002-Ab635p, NanoTag), anti-mouse FluoTagX2 IgG1 565 (#N2702-At565s, NanoTag), anti-mouse FluoTagX2 IgG2a/b 635p (#N2702-Ab635p-S,, NanoTag), anti-rabbit FluoTagX4 580 (#N2404-Ab580-S, NanoTag), anti-rabbit FluoTagX4 635p (#N2404-Ab635p, NanoTag), goat anti-chicken 488 (#A11039, Invitrogen), goat anti-guinea pig 488 (#A-11073, ThermoFisher Scientific), goat anti-mouse IgG STAR580 (#ST580-1001, Abberior), goat anti-rabbit STAR635p (#ST635P-1002, Abberior), TO-PRO^TM^-3 Iodide 642/661 (#T3605, ThermoFisher Scientific). Depending on the specific acquisition system used, samples were then mounted using either ProLongTM Glass Antifade Mountant (#P36984, Life Technologies Corporation) or Mowiol mounting medium (#0713, Carl Roth GmbH + Co. KG).

### Palm3 antibodies

As immunogens, two mouse Palm3 sequences were selected that are non-overlapping, and do not contain sequence stretches with similarity to the other paralemmin isoforms (i.e., they exclude the N-terminal coiled-coil region and the C-terminal CaaX motif): Palm3-p31 (aa 114-399; QSAST…GEASWE) and Palm3-p32 (aa 404-725; VEKVE…ASGPK). These inserts were amplified by RT-PCR from mouse brain RNA, cloned into the SmaI site of pQE-82L, and verified by sequencing (RefSeq XP_486140). His-tagged fusion proteins were expressed in bacteria and affinity-purified on nickel-agarose. As both fusion proteins were partially insoluble, affinity purifications were performed on the native, soluble fractions as well as on the denatured, urea-solubilized fractions, following Qiagen protocols. Rabbits were immunized separately with native and denatured immunogens at Pineda-Abservice (Berlin, Germany), and sera were affinity-purified on the respective CNBr- or tresyl-agarose-coupled immunogen proteins. Antibody preparations anti-Palm3 p31 rb2 nat (fr2-3), anti-Palm3 p32 rb1 den, and anti-Palm3 p31 rb1 den (fr2-3) were employed in the present study as indicated.

### Lightsheet microscopy

Cytocochleograms were performed in hearing animals at p21. Following overnight fixation in 4% FA in PBS at 4°C, osseous labyrinths were decalcified at RT for four days in 10% EDTA. Chemical tissue clearing and immunolabeling were performed using a modified version of the iDISCO+ protocol (Renier *et al*, 2016; Rankovic *et al*, 2021).

All incubation and washing steps were performed on a thermomixer at 37°C and at RT, respectively. After permeabilization and blocking, a mixture of the primary antibody rabbit anti-Myosin7a (#PTS-25-6790-C050, Enzo) and nanobody anti-rabbit FluoTagX4 IgG 635p (#N2404-Ab635p, NanoTag) was prepared and preincubated for 45 min at RT. The primary antibody/nanobody mixture was then diluted in blocking solution and applied to the samples for 6 days. After washing, samples were cleared in 66% dichloromethane (#270997, Sigma-Aldrich)/33% methanol for 3 h at RT, followed by 2 x 15 min incubation in 100% dichloromethane, and stored in dibenzyl ether (#33630, Sigma-Aldrich). Imaging was performed using a LaVision UltraMicroscope II (NA 0.154, sheet thickness 5 µm) with 3 µm step size and 8-10x total zoom (2x objective, 4-5x optical zoom). OHC survival along the tonotopic axis was analyzed using a custom MATLAB script (Rankovic *et al*, 2021) implementing Greenwood’s function (Greenwood, 1961) to map frequency regions (4, 8, 12, 16, 24 and 32 kHz) based on mouse cochlear parameters (Müller *et al*, 2005). OHCs were manually counted using the Cell Counter tool in ImageJ/Fiji (Schneider *et al*, 2012) in respect to their approximate frequency regions.

### OHC length and fluorescence intensity analysis

OHC lengths were measured using the straight-line tool in ImageJ/Fiji (Schneider *et al*, 2012). Fluorescence intensities of Palm3, Prestin, and ɑII-Spectrin were quantified from maximal projections of sample images in ImageJ/Fiji (Schneider *et al*, 2012). Background fluorescence was measured in a 3 µm x 3 µm area lacking apparent fluorescence intensity, subtracted from the image, and average intensities within 50 µm x 50 µm regions containing OHCs were then measured. Average intensity values were normalized to control.

To quantify spectrin fluorescence intensities in WT, *Palm3*-KO, and *Palm3*-rescue animals, OHC masks were generated in *Imaris* 10.2.0 using the *cells* algorithm, based on Myosin7a staining (filter width: 0.8 µm; largest sphere diameter: 3 µm). Masks were expanded to include membranes, and adjacent cells were split using a 6.0 µm diameter threshold. The resulting 3D reconstructions were exported to *surfaces* for average fluorescence intensity analysis. Cuticular plate intensities were analyzed by creating 3D *surfaces* renderings from spectrin staining (surface detail: 0.3 µm; largest sphere diameter: 0.4 µm). All average intensity values were normalized to controls.

### Patch-clamp electrophysiology on OHCs

Animal care and tissue collection from *Palm3*-KO and wildtype mice followed federal and institutional guidelines at the Philipps University Marburg. Mice of the indicated age were killed by decapitation under isoflurane anesthesia. Cochleae were dissected on ice in a solution containing (in mM) 144 NaCl, 5.8 KCl, 1.3 CaCl_2_, 0.9 MgCl_2_, 10 HEPES, 0.7 Na_2_HPO_4_, 5.6 glucose (pH adjusted to 7.4 with NaOH). Recordings were done within 2 hours after dissection. For patch-clamp recordings, pieces of the apical cochlear turn were mounted in a recording chamber with a flexible glass fiber glued to the bottom coverslip at one end. The bath chamber was continuously perfused with the same extracellular solution used for dissection. For establishing whole-cell patch-clamp recordings, the preparation was viewed with an upright BX51WI microscope equipped with water immersion objectives (Olympus, Tokyo, Japan). Whole-cell patch-clamp recordings were performed with EPC10 amplifiers (Heka, Lambrecht, Germany) controlled by Patchmaster software (Heka). Patch pipettes were pulled from borosilicate glass to obtain open tip resistances of 2-4 MOhm when filled with intracellular solution. All experiments were performed at room temperature (21-23°C).

### Non-linear capacitance measurements

Recording pipettes were filled with intracellular solution containing (mM): 100 CsCl, 20 TEA-Cl, 3.5 MgCl_2_, 0.1 CaCl_2_, 6 BAPTA, 2.5 Na_2_ATP, 5 HEPES (pH adjusted to 7.4 with NaOH, Osmolarity adjusted to 300 mOsm/kg with glucose). Whole-cell series resistance was < 10 MOhm. NLC was measured using a stair-step protocol as described (Oliver *et al*, 1998; Huang & Santos-Sacchi, 1993), with a step duration of 10 ms at voltage increments of 10 mV. Currents were low-pass filtered at 10 kHz and sampled at 100 kHz. In these experiments, whole-cell membrane capacitance (C_M_) was not compensated. The time constant of current decay (t) was determined from mono-exponential fits to the current transients, and input resistance (Rin) was calculated from steady-state currents at the end of each voltage step. Integration of the current transients yielded the charge (Q). Cm(i) at each incremental voltage step i was derived according to (Huang & Santos-Sacchi, 1993), taking into account the relatively low Rm of OHCs:

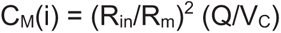

and

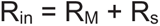

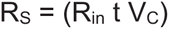

RS is the series resistance; RM is the membrane resistance and VC is the command voltage step amplitude (10 mV). CM (i) was plotted versus (V(i) + V(i + 1))/2, after correction of all voltages for series resistance errors.

Voltage-dependent capacitance was fitted with a derivative of the Boltzmann function:

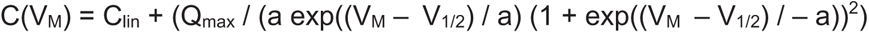

where Clin is the linear membrane capacitance, VM is the membrane potential, Qmax is the maximum voltage sensor charge moved through the membrane electrical field, V1/2 is the voltage at half-maximal charge transfer, and a is the slope factor of the voltage-dependence. Clin reflects the surface membrane area of OHCs. As a measure for prestin density in the OHC membrane, NLC is also presented normalized to Clin.

### Polymer resin (melamine) embedding

Melamine was prepared following the protocol described in (Revelo *et al*, 2014). Briefly, 48 mg of *p*-toluenesulfonic acid monohydrate (#402885, Sigma Aldrich) was dissolved in 0.576 mL distilled water, followed by the addition of 1.344 g 2,4,6-Tris[bis(methoxymethyl)amino]-1,3,5-triazine (#T2059, TCI chemicals) and vortexed for 2 h until completely dissolved. After fixation and immunostaining, each organ of Corti was placed into the head of a block mold (#E69931, Easy-Molds™) and 100 µL of freshly prepared melamine was poured inside the mold until the organ of Corti was covered. To facilitate melamine penetration into the tissue, samples were incubated overnight at RT in a vacuum desiccator. Melamine was polymerized at 40°C for 24 h. Subsequently, the block mold was filled up with Epoxy resin (#40200029, EpoFix kit, Struers) and cured at 60°C for 48 h. Melamine blocks were sectioned into ∽80-90 nm ultra-thin sections using a Reichert Ultracut S microtome (Leica Microsystems, Vienna, Austria) with a diamond knife (Diatome Ultra 45°, Switzerland, Nidau) at 3-6° clearance angle. Sections were mounted in Mowiol for 2D stimulated emission depletion (STED) microscopy.

### Li’s intensity correlation quotient (ICQ) and nearest neighbor distance (NND) analysis

STED images were pre-processed to subtract background fluorescence in ImageJ/Fiji software (Schneider *et al*, 2012). A 3 µm x 3 µm area lacking apparent fluorescence intensity was measured to determine average background intensity, which was then subtracted from the image. ICQ values obtained using Li’s intensity correlation analysis (ICA; Li *et al*, 2004) using the Colocalization Analyzer on Huygens Professional software (Scientific Volume Imaging, Hilversum, The Netherlands). Nearest neighbor distances (NND) of Palm3 immunolabeling were analyzed in *Imaris* 10.2.0 (Oxford Instruments) using the spot-detection algorithm with an estimated XY diameter of 0.08 µm. Thresholding based on the average Palm3 signal intensity was applied to remove noise, and the few detected spots that were not localized within the cells of interest were manually excluded. NND was calculated from the shortest distance between the centers of homogeneous masses of detected spots. Figure 6A was deconvolved for representation purposes with Huygens Professional version 25.04 using the “Standard” Deconvolution Express Profile with Acuity set to: −5.3 (Scientific Volume Imaging, The Netherlands).

### High pressure freezing and freeze-substitution

High pressure freezing (HPF) and freeze-substitution (FS) and embedding in epoxy resin (EPON) were essentially carried out as described in (Wong *et al*, 2014; Chakrabarti *et al*, 2018, 2022). For HPF, apical organs of Corti were placed within a type A aluminum specimen carrier (Leica Microsystems, Wetzlar, Germany) with a depth of 0.2 mm, filled with HEPES-HBSS NaCl 141.7, HEPES 10, KCl 5.36, MgCl2 1, MgSO4*7H2O 0.5, freshly added: glucose 2 mg/ml and L-glutamine 0.5 mg/ml (pH adjusted to 7.2 with NaOH. Osmolarity ≤ 300 mmol/kg). A type B carrier was dipped into 1-hexadecene (Sigma-Aldrich, Darmstadt, Germany) and positioned adjacent to the sample. The samples were promptly frozen using the HPM100 (Leica Microsystems, Wetzlar, Germany) and directly placed in liquid nitrogen for storage. For FS, the EM AFS2 (Leica Microsystems, Wetzlar, Germany) was utilized. Samples were incubated in 0.1% (w/v) tannic acid in acetone for four days at −90°C. Samples were washed three times with acetone for 1 h each. Thereafter, 2 % (w/v) osmium tetroxide in acetone was added and incubated for 7 h. The temperature was increased in incremental steps of 5°C per hour to −20°C. At −20°C, samples were incubated for another 17 h before the temperature was raised to 4°C in 10°C increments per hour. While the samples were being heated to room temperature, they were washed on three occasions with acetone, each for 1 h. Following the washing process, the samples were infiltrated with 1:1 EPON (Agar pre-mix kit, Plano GmbH, Wetzlar, Germany) in acetone solution and incubated for 3-6 h. The solution was replaced with 100% EPON and incubated overnight. Then, EPON was replaced with fresh EPON and incubated for an additional 3-6 h. All incubation steps were conducted on a rotary wheel at RT. Subsequently, the samples were positioned within embedding molds (Agar Scientific Ltd., Stansted, Essex, United Kingdom), filled with 100% fresh EPON, and polymerized in an incubator (Incu-line, VWR, Darmstadt, Germany) set to 70°C for 48 h.

Samples that did not exhibit the desired orientation following embedding were reoriented in order to allow for the acquisition of a cross-section through the organ of Corti (in the region of the apical 1.5 turns). The portion of the EPON containing the sample was sectioned with a saw, placed in embedding molds (Agar Scientific Ltd., Stansted, Essex, United Kingdom) oriented as desired, filled with EPON, and polymerized for 48 h at 70°C in the drying oven Incu-line (VWR, Darmstadt, Germany). For this purpose, the AGR1140 Agar 100 Premix Kit – Hard (Agar Scientific, London, United Kingdom) was used. The EPON directly surrounding the sample was roughly trimmed using a file (Titania Solingen GmbH, Wülfrath, Germany) and a razor blade (Plano GmbH, Wetzlar, Germany) into the shape of a trapezoid and subsequently smoothened at the block face with the Cyrotrim 45° (DiATOME, Nidau, Switzerland) in the EM UC7 ultramicrotome (Leica Microsystems, Wetzlar, Germany).

Following the trimming of the samples, 75 nm ultrathin sections were obtained for transmission electron microscopy (TEM) analysis of the immunogold label and 2D analyses. This was achieved by approaching the block face with a 35° diamond knife (DiATOME, Nidau, Switzerland) using the EM UC7 ultramicrotome (Leica Microsystems, Wetzlar, Germany). For each sample, a total of 3-5 sections were applied onto 17-45 1 x 2 mm copper slot grids (3.05 mm Ø; Plano, Germany) coated with 1% formvar (w/v in water free chloroform). For electron tomography (ET), 250 nm semithin sections were generated and 3-5 sections were applied to 20-35 75 copper mesh grids (3.05 mm Ø; Plano, Germany), which were coated with 1% formvar.

Both the grids for TEM as well as for ET analysis were stained by incubating them on droplets of Uranyless (lanthanum, gadolinium and dysprosium salts in distilled water, Delta Microscopies, Mauressac, France) for 20 to 25 minutes. Subsequently, the grids were washed six times on droplets of distilled water, for one minute each. For ET samples, droplets of 10 nm gold nanoparticles (10 nm gold colloid, BBI solutions, Freiburg, Germany) were applied to the mesh grids for ET by incubating from both sides for 8 minutes each. The gold nanoparticles serve as fiducials for tomogram alignment.

### Transmission electron microscopy

TEM examination for comparison of the abundance of SSC employed the JEM 1011 TEM (JEOL, Freising, Germany). For each sample one to four grids were examined and 2D micrographs of OHC were obtained at different magnifications. Overview micrographs of the examined cochlear region were taken at 800-fold magnification to determine whether the cells are indeed OHC. Micrographs of the individual OHC were obtained at 2,500-fold magnification and close-ups of the plasma membrane of each OHC in the region of nucleus at 6,000-fold magnification.

Only OHCs that were unambiguously identified as such, based on their localization and morphology, were examined. Therefore, stereocilia had to be in close proximity to the cells and the tunnel of Corti as well as the IHC and cochlear nerve fibers had to lie medial to the cells. Since the region of interest (ROI) was the plasma membrane in the region lateral to the nucleus, only OHC with a visible nucleus were imaged. The final criterion was the integrity of the plasma membrane to ensure a near-native state of the OHC.

### Electron tomography

Single tilt series of the area lateral to the nucleus of the OHC was acquired using the Serial-EM software package (Mastronade, University of Colorado, USA) (Kremer *et al*, 1996a) at a JEM 2100Plus TEM (JEOL, Freisingen, Germany) essentially performed as described in (Hintze *et al*, 2024). This TEM is equipped with a Xarosa camera (EMSIS GmbH, Münster, Germany) and the RADIUS software package (EMSIS GmbH, Münster, Germany), used for finding the ROI, and operates at 200 kV. The tilt series, if possible, spanned a range of 1° increments from −55 to +55° and were obtained at either 12,000x or 20,000x magnification. Subsequently, tomograms of the OHC were generated using the IMOD software package etomo (Mastronade, University of Colorado, USA) (Kremer *et al*, 1996b). In this process, one pixel corresponded to a distance of 1.758 nm at a 12,000-fold magnification and of 1 nm at 20,000-fold magnification.

### Analysis of SSC-TEM data

2D micrographs obtained via TEM at 6,000-fold magnification were used to quantify alterations in the number of SSC between WT and *Palm3*-KO mice. For this purpose, the plasma membrane lateral to the nucleus of each OHC was traced and the length was measured using the “freehand line” tool in the Fiji ImageJ software (Schindelin *et al*, 2012). Pixels were converted into nanometers by measuring the scale bar in 6000x magnified micrographs using the straight-line tool to determine a conversion factor, with one nanometer corresponding to approximately 0.95 pixels. To determine the number of SSC per micrometer of plasma membrane, the SSC in the ROI of each OHC were enumerated.

### Analysis of SSC-ET data

Distinguishable SSC on each virtual section of each tomogram were sculpted manually on their membrane, using the “sculpting” tool of the IMOD 3dmod software package etomo (Mastronade, University of Colorado, USA) (Kremer *et al*, 1996b). The SSC belonging to a single tomogram were treated as a closed object and meshed into a 3D model without a cap. For this object, the overall surface area of the mesh, as well as the total volume of the outline and the volume within the mesh, were determined.

In order to put the position of the SSC within the OHC into perspective, the plasma membrane of the OHC lateral to the nucleus and the mitochondria, when present, were incorporated into the 3D models. The plasma membrane and the mitochondria were manually sculpted on every third virtual section and transferred to the sections in between using the “interpolator” tool in IMOD 3dmod (Mastronade, University of Colorado, USA) (Kremer *et al*, 1996b), provided that their course exhibited only slight variations. Otherwise, the procedure described for the SSC was employed.

To facilitate quantitative comparison of the localization of the SSC between the WT and *Palm3*-KO mice, the shortest distance between the outer SSC membrane and the inner border of the plasma membrane of the OHC was determined on the virtual sections of the tomograms using the “measuring” tool in IMOD 3dmod (Mastronade, University of Colorado, USA) (Kremer *et al*, 1996b). Consistency and reliability of the results was ensured, by measuring this distance between the SSC and the plasma membrane nine times for each tomogram at different positions. The initial three measurements were obtained from the virtual section that marked the midpoint of the entire series. In this instance, the shortest distance was determined for the SSC situated at the middle of the section as well as for the SSC located at either ends. Measurements were repeated on the 50^th^ section above and below the central section for the tomograms obtained at 20,000x magnification and on the 20^th^ section above and below for the tomograms obtained at 12,000x magnification.

### rAAV expression construct and viral vector production

A fusion cDNA encoding yellow fluorescent protein (YFP) and mouse Paralemmin-3 (from codon 2 [ALQTP…] to the stop codon), connected by the 20 aa linker GSSPSTSLYKKAGSAAAPFT, was originally cloned into Invitrogen Vivid Colors pcDNA6.2/N-YFP-DEST. This fusion cDNA was subcloned, by InFusion cloning (TAKARA) according to the manufacturer’s instructions, into a pAAV backbone, between the human *Myo15a* promotor and a non-oncogenic variant of WPRE (Zanta-Boussif *et al*, 2009) and the bovine growth hormone derived poly-adenylation signal. Sequence integrity was confirmed by Sanger sequencing. This vector was amplified in recombinase deficient *E. coli* NEB Stable (NEB) and purified using the NucleoBond PC 2000 EF Mega kit for endotoxin free plasmid DNA (Macherey-Nagel) according to the manufacturer’s instructions.

Viral vector purification was performed essentially as previously published (Keppeler *et al*, 2018) and an extensive description is available in (Huet & Rankovic, 2021). In brief, triple transfection of HEK-293T cells was performed using the pHelper plasmid (TaKaRa, USA), trans-plasmid with the AAV-S capsid (Ivanchenko *et al*, 2021) and cis-plasmid with *YFP-Palm3* under the control of the human *Myo15a* promotor. Cells were regularly tested for mycoplasma contamination. Viral particles were harvested 72 hours after transfection from the medium and 120 h after transfection from cells and the medium. Viral particles from medium were precipitated with 40% polyethylene glycol 8000 (Acros Organics, Germany) in 500 mM NaCl for 2 h at 4 °C and then after centrifugation combined with cell pellets for processing. Pellets were suspended in 500 mM NaCl, 40 mM Tris, 2.5 mM MgCl2, pH 8, and 100 U/ml of salt-activated nuclease (Arcticzymes, USA) at 37 °C for 2h (cells) or 30 min (medium). Afterward, lysates were clarified by centrifugation at 2000 ×g for 10 min and viral vectors were purified over iodixanol (Optiprep, Axis Shield, Norway) step gradients (15%, 25%, 40%, and 60%) at 350,000 × g for 2.25 h. Viral vectors were concentrated using Amicon filters (EMD, UFC910024) and formulated in sterile phosphate-buffered saline (PBS) supplemented with 0.001% Pluronic F-68 (Gibco, Germany). Viral vector titer was determined by the number of DNase I resistant vg using AAV titration kit (TaKaRa/Clontech) with adaptations to usage in crystal digital PCR (dPCR). Briefly, dilutions of purified DNA were subjected to crystal dPCR using 5x Naica PCR reaction mix (Stilla, R10056) supplemented with primers provided in the AAV titration kit, 0.8 mg/µl Alexa 647 (Invitrogen, 11570266) and 1.5x Eva Green (VWR, 31077-T). Formation of droplet crystals and PCR was performed in a Naica Geode (Stilla) using Naica Ruby or Sapphire Chips (Stilla) and finally analyzed in the Naica Prism3 reader (Stilla) equipped with Crystal Reader and Crystal Miner softwares (Stilla). Purity of produced viruses was checked by silver staining (Pierce, Germany) after gel electrophoresis (Novex™ 4–12% Tris–Glycine, Thermo Fisher Scientific) according to the manufacturer’s instruction. Viral stocks were kept at −80°C until usage.

### Postnatal AAV-Injection

As systemic analgesia, *Palm3*^-/-^ pups received subcutaneous injections of Buprenorphine (0.1 mg/kg) and Carprofen (5 mg/kg) 30 min prior to the AAV-injection procedure. During the procedure, anesthesia was induced with 5% isoflurane and maintained at 3% isoflurane. *AAV*-S.*YFP-Palm3* was injected into the left ear via the round window method (Jung *et al*, 2015) at postnatal day 4-6. Hearing assessment (ABR and DPOAE measurements) and cochlear morphology were assessed 4-5 weeks post-injection. The virus solution was injected in a concentration of 2.66×10^12^ genome copies/ml. All virus injection experiments were performed in compliance with the national animal care guidelines and were approved by both the board for animal welfare of the University Medical Center Göttingen and the State Animal Welfare office of Lower Saxony (AZ19/3133).

### Confocal and STED microscopy

Confocal imaging was performed using either a Leica SP8 microscope (Leica Microsystems, Wetzlar Germany) or an Olympus-IX83 based Abberior Instruments Expert Line 775 nm 2-color STED microscope (Abberior Instruments GmbH), operated in confocal mode. 2D-STED images were acquired on a Nikon Eclipse Ti2-E based (Nikon, Tokyo, Japan) STEDYCON system (Abberior Instruments GmbH) with a pulsed 775 nm depletion laser. Lasers with wavelengths of 488, 587, and 633 nm were used for excitation. All 2D-STED images were acquired using a 100x oil immersion objective (1.45 NA), with a final resolution of 54-56 nm.

### Prestin average fluorescence intensity

Prestin average intensity analysis was performed by generating a mask of OHCs using the *surfaces* algorithm in *Imaris* 10.2.0. For non-transduced OHCs, 3D reconstructions of Prestin immunofluorescence were generated (surface detail: 0.15 µm, largest sphere diameter: 0.6 µm). For YFP-Palm3-transduced OHCs, YFP-Palm3 immunofluorescence signal was rendered (surface detail: 0.163 µm, largest sphere diameter: 0.611 µm). Prestin average intensity values from YFP-Palm3*-*transduced OHCs were normalized to values from non-transduced controls. Prestin coefficient of variation analysis was performed using the ImageJ/Fiji. Maximum projections were background-corrected by subtracting intensity measured in a 3 µm x 3 µm area lacking apparent fluorescence intensity. Mean and standard deviation were calculated from a 3 µm x 3 µm region within an OHC in the Prestin fluorescence channel by plotting a histogram. The coefficient of variation was determined by dividing the standard deviation by the mean.

### Statistics

Statistical analyses were conducted using Prism 10 (GraphPad Software Inc, La Jolla, CA). Normality of the data distribution was assessed with Kolmogorov-Smirnov normality test. For comparisons between two groups, the two-tailed Wilcoxon matched-pairs signed-rank test was used for paired data (e.g., IHC rescue cell count), and the two-tailed Mann-Whitney test for unpaired, non-normally distributed data (e.g., spectrin intensity in WT vs *Palm3*-KO). Unpaired multiple group comparisons (e.g., OHC degeneration along the tonotopic axis), were analyzed with two-way ANOVA and Holm-Šídák’s correction. For paired multiple comparisons (e.g., rescue experiments), two-way repeated measures ANOVA with Holm-Šídák’s correction was applied. Single value plots display individual data points with mean ± standard error of the mean (SEM). In violin plots, medians are indicated by solid lines and quartiles with dashed lines. *N* indicates the number of animals, while *n* indicates the number of ears or hair cells, as specified accordingly in figure legends. Statistical significance is indicated by the symbols *n.s.* for not significant, \**P* < 0.05, \*\**P* < 0.01, \*\*\**P* < 0.001, and \*\*\*\**P* < 0.0001. A comprehensive summary table with all statistical information can be found in the Supplement of this article (**Supplemental Tables 1-3**).

## Results

### Genetic deletion of *Palm3* results in severe, early onset and progressive hearing loss in mice

To expand the initial findings of the GMC phenotypic screen, we extended the analysis window for the auditory brainstem response (ABR) and distortion product otoacoustic emission (DPOAE) measurements to juvenile (3-week-old) and young-adult (12-13-week-old) *Palm3*-KO mice, along with their respective age-matched controls. We found an age-dependent reduction in ABR Wave I (**Figure 1A-A’**). ABR thresholds to pure tone and click stimulation were statistically significantly increased by 15-30 dB in juvenile *Palm3*-KO mice and then further deteriorated in the older age group, where thresholds were elevated by 68 dB for clicks and mostly exceeded our loudspeaker limit of 80 dB SPL for tone bursts (**Figure 1B*-*B’**). To additionally evaluate OHC-mediated active cochlear amplification, distortion-product otoacoustic emissions (DPOAEs) were recorded and found to already be markedly diminished within the young age group (**Figure 1C-C’**), with a progressive decline leading to near-complete loss of DPOAEs by 12-13 weeks of age (**Figure 1D*-*D’**). This latter finding is strongly indicative of corrupted OHC function or cell loss.

**Figure 1.**
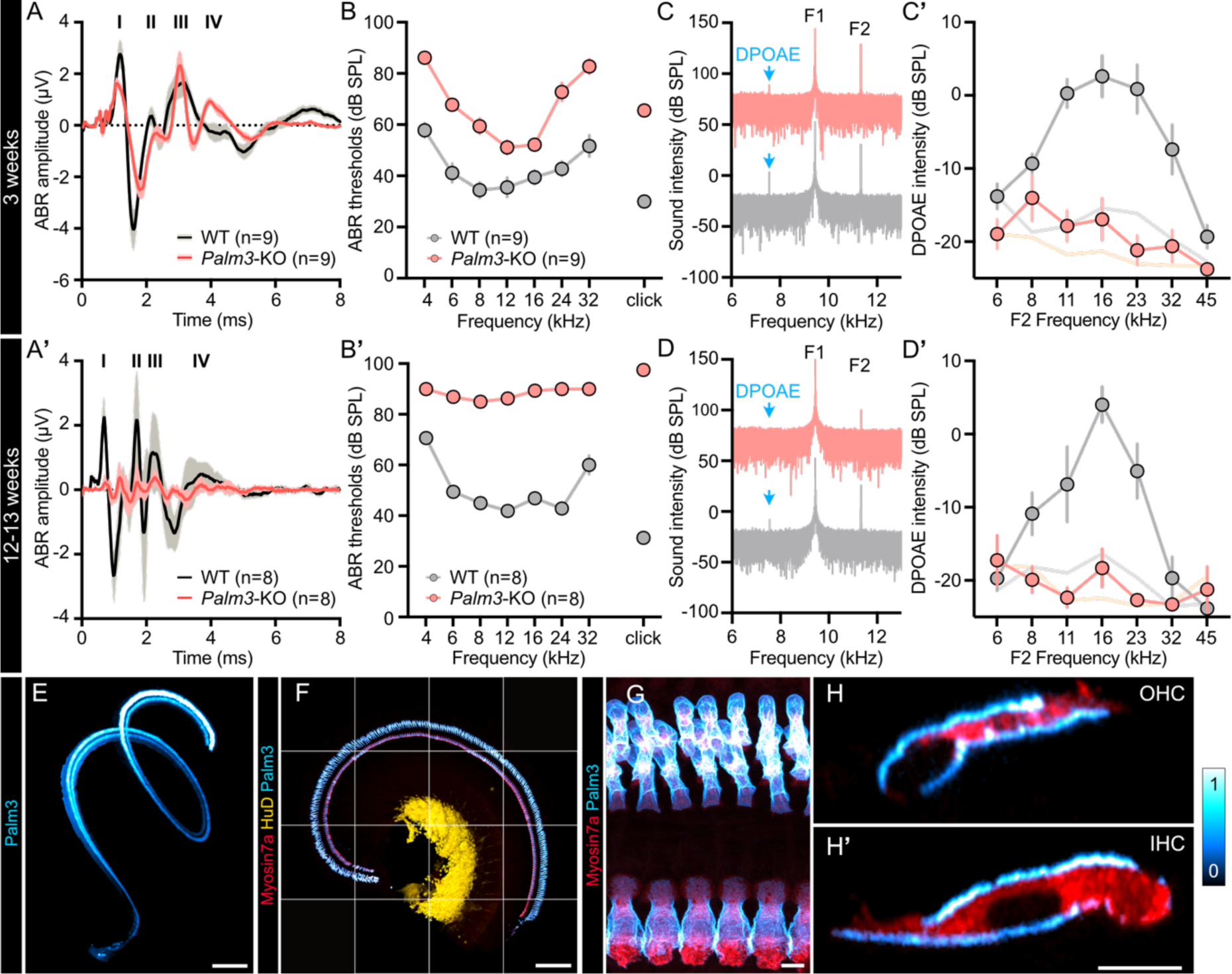
Palm3-KO mice exhibit early onset and progressive hearing loss. (A,A’) Grand averages of auditory brainstem response (ABR) waveforms upon 100 dB SPL click stimulation at 20 Hz showed (A) a moderate amplitude reduction in 3-week-old and (A’) a stark reduction in 12-13-week-old *Palm3*-KO mice compared to WT-littermates. (B,B’) ABR threshold measurements from (B) 3-week-(*N*_animals_= 9 for each WT and *Palm3-*KO) and (B’) 12-13-week-old (*N*_animals_= 8 for each WT and *Palm3-*KO) WT (grey) and *Palm3*-KO (pink) mice revealed early-onset and progressive hearing impairment in *Palm3*-KO mice. Tone burst threshold values that surpassed the maximum output of the loudspeaker of 80 dB SPL were assigned as 90 dB for the purpose of calculating means ± SEM. (C,D) Representative frequency spectra of DPOAE recordings for the specified age groups for the stimulus combination F1 9.4 kHz 60 dB, F2 11.3 kHz 50 dB, expected DPOAE at 7.5kHz. *Palm3*-KO traces are offset by +100 dB to avoid overlap. Note the complete absence of a detectable DPOAE in the 12-13-week-old cohort. (C’,D’) DPOAE amplitude in response to paired simultaneous sine waves in (C’) 3-week-(*N*_animals_= 9 for WT and *N*_animals_= 8 for *Palm3-*KO) and (D’) 12-13-week-old (*N*_animals_= 7 for each WT and *Palm3-*KO) *Palm3*-KO (pink) mice are attenuated compared to WT littermates (grey). Light grey and light orange lines indicate the average noise floor. (E) Representative maximum z-projection of lightsheet microscopy on iDisco-cleared cochlea of a P14 WT mouse. Hair cells are stained with Palm3 (cyan hot). Lightsheet microscopy image was processed as described in **Supplementary figure 2**. Scale bar: 100 µm. (F) Representative maximum z-projection showing an overview of the distribution of Palm3 expression in the organ of Corti from a P16 WT mouse. Hair cells are immunostained for Palm3 (cyan hot) and Myosin7A (red), SGN somata are immunostained for HuD (yellow). Scale bar: 150 µm. (G) Localization of Palm3 to the lateral plasma membrane of auditory hair cells at 100X magnification, assessed in the apical cochlear turns of a P16 WT mouse. Scale bar: 5 µm. (H-H’) Representative yz-projections of single optical sections of an OHC and an IHC. Scale bar: 5 µm. Data information: Comparison of genotypes: Two-way ANOVA with Holm-Šídák’s multiple comparisons correction: \*\*\*\**P* <0.0001; Mean ± SEM.

### In the murine cochlea, Palm3 is specifically and exclusively expressed in auditory hair cells

To determine the expression of Palm3 within the peripheral auditory system, we conducted immunohistochemical stainings on acutely dissected organs of Corti from post-hearing onset wildtype mice. Samples were labeled using antibodies against Palm3, the cytosolic hair cell marker Myosin7a, and the neuronal soma marker HuD (**Figure 1F**). In these experiments, Palm3 localized specifically and exclusively to the lateral plasma membrane of both OHCs and IHCs (**Figure 1G-H’**). To verify target specificity, we employed three independent antibodies raised and affinity-purified against two distinct, non-overlapping and Palm3-specific immunogen sequences: P31 and P32 (see Methods). All three antibodies produced identical staining patterns, labeling only the lateral plasma membranes of both OHCs and IHCs, with no detectable signals in other cell types within the sensory epithelium or the spiral ganglion and a complete loss of staining in *Palm3*-KO preparations (**Supplementary figure 1**). This cell-type specific expression aligns well with the publicly available single-cell RNA sequencing data of auditory hair cells deposited on gEAR (**Supplementary figure 3**; Orvis *et al*, 2021; https://umgear.org/). Palm3 was also detected in the lateral membranes of hair cells within the vestibular system – including the macular organs and *cristae ampullaris* (**Supplementary figure 4**).

### Hearing loss in *Palm3*-KOs results from progressive hair cell degeneration

Given the hearing deficits observed in *Palm3*-KO mice, which point toward dysfunction within the peripheral auditory pathway, and the specific localization of Palm3 to the lateral plasma membranes of IHCs and OHCs, we sought to elucidate the underlying mechanism by first assessing the fate of IHCs and OHCs in the apical cochlear turn of *Palm3*-KO mice over a time course from 2 to 50 weeks of age. Using Myosin7a as a marker for hair cells and the DNA marker ToPro3 to label all nuclei for structural context, we found a pronounced and progressive loss of OHCs starting as early as 2 weeks of age, with degeneration worsening up to 50 weeks of age, where OHCs were almost completely lost (**Figure 2A-B**). These results are consistent with the severely reduced DPOAE amplitudes in *Palm3*-KO mice (**Figure 1C-D’**). Additionally, a mild but statistically significant loss of IHCs was observed (**Figure 2B**). Notably, regardless of the specific subpopulation, remaining *Palm3*-KO hair cells displayed a more compact and aberrant morphology in comparison to their age-matched WT counterparts (**Figure 3A, B, C, C’**).

**Figure 2.**
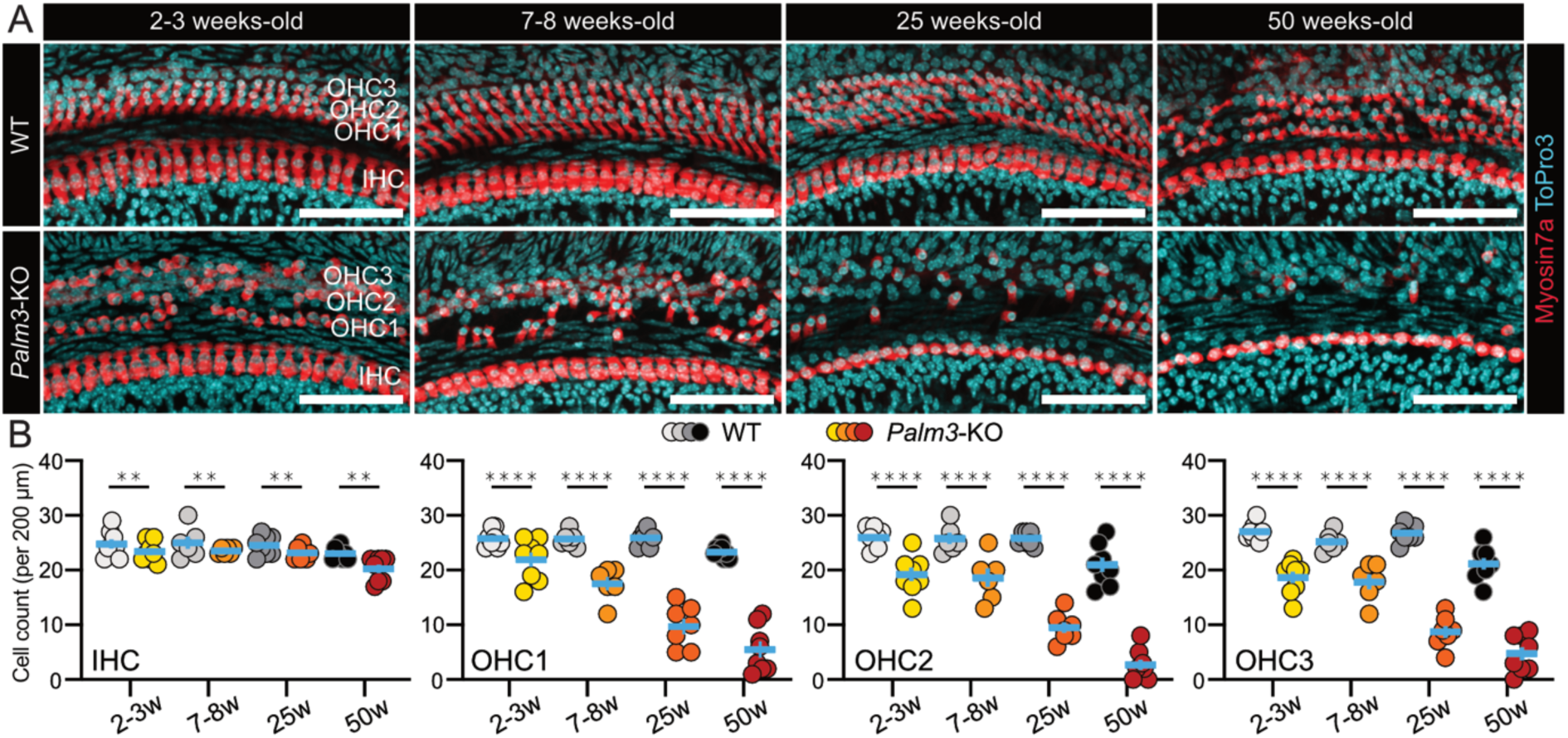
Hearing loss in Palm3-KO is due to progressive hair cell degeneration. Hair cell survival was assessed in the apical cochlear turn (8-11 kHz frequency range) of 2-3 week-, 7-8 week-, 25 week- and 50-week-old WT (upper panels) and *Palm3*-KO animals (lower panels). Organs of Corti explants were immunostained against the hair cell marker Myosin7a (red) and the DNA stain ToPro3 (cyan). Scale bars: 50 μm. (A) Quantitative assessment of hair cell counts in whole-mount apical organs of Corti revealed *Palm3*-KO mice show a small decrease in the survival rate of IHCs, whereas OHCs show varying degrees of progressive degeneration in respect to the different rows. 2-3 weeks old: *N*_animals_= 4, *n*_Corti_ = 8 for each WT and *Palm3*-KO, 7-8 weeks old: *N*_animals_= 3, *n*_Corti_ = 6 for each WT and *Palm3*-KO, 25 weeks old*: N*_animals_= 4, *n*_Corti_ = 8 for WT and *N*_animals_= 4, *n*_Corti_ = 7 for *Palm3*-KO, 50 weeks old: *N*_animals_= 4, *n*_Corti_ = 8 for each WT and *Palm3*-KO. Data information: Two-way ANOVA with Holm-Šídák‘s multiple comparisons correction: \*\**P* <0.01; \*\*\*\**P* <0.0001; Mean ± SEM.

**Figure 3.**
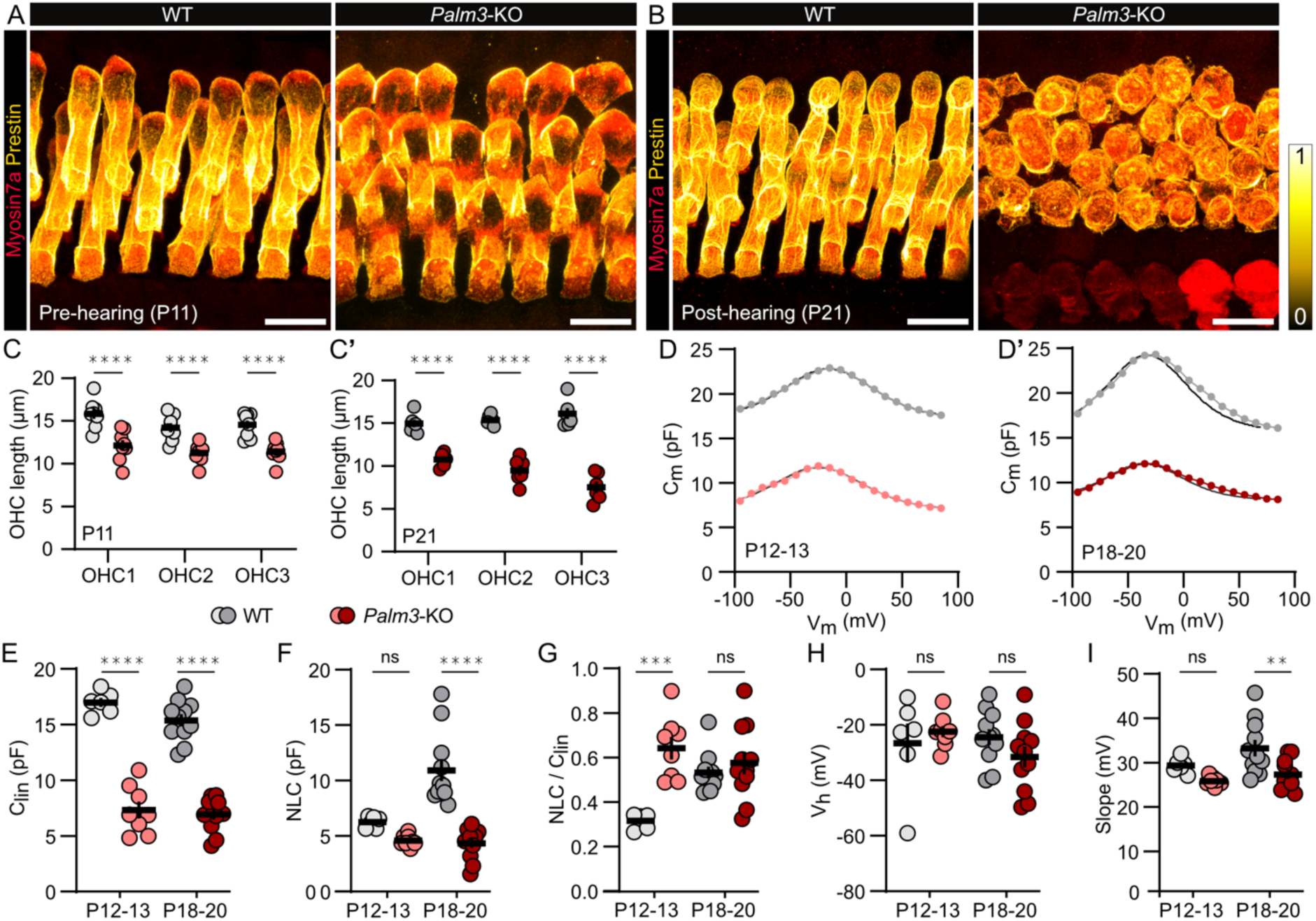
Genetic loss of Palm3 resulted in reduced OHC length, perturbed localization of Prestin and decreased electromotile capacity. (A,B) Representative confocal z-y projections of apical organs of Corti from WT and *Palm3*-KO at (A) P11 and (B) P21. OHCs were immunostained against the hair cell marker Myosin7a (red) and Prestin (yellow hot). Scale bars: 10 µm. (C,C’) *Palm3*-KO organs of Corti reveal a dramatic reduction in OHC lengths regardless of row position at (C) P11 (*N*_animals_= 4, *n*_Corti_ = 8 for WT and *N*_animals_= 4, *n*_Corti_ = 8 for *Palm3*-KO), which progressed until (C’) P21 (*N*_animals_= 3, *n*_Corti_ = 6 for WT and *N*_animals_= 3, *n*_Corti_ = 6 for *Palm3*-KO). (D-D’) Example traces of voltage-dependent cell capacitance from (D) P12-13 and (D’) P18-20 WT and *Palm3-*KO OHCs measured with the stair-step stimulus protocol from −100 mV to 85 mV in 10-mV increments and 5 ms step duration. (E-I) Quantitative analysis derived from (D,D’) shows *Palm3*-KO OHCs to have (E) an overall decrease in linear capacitance (C_lin_), (F) reduced non-linear capacitance (NLC) at P18-20, (G) increased ratio of NLC to C_lin_ (NLC/ C_lin_) at P12-13, (H) unchanged voltage at peak capacitance (V_h_), and (I) decreased slope at P18-20. P12-13: *n*_cells_ = 6 for WT and *n*_cells_ = 8 for *Palm3*-KO; P18-20: *n*_cells_ = 11 for WT and *n*_cells_ = 9 for *Palm3*-KO. Data information: Two-way ANOVA with Holm-Šídák‘s multiple comparisons correction: ns = not significant; ** *P* < 0.01; *** *P* < 0.001 **** *P* < 0.0001; Mean ± SEM.

To test for tonotopic differences, we acquired *in situ* cytocochleograms to examine frequency-specific OHC degeneration in *Palm3*-KO mice, using a modified iDisco protocol (Keppeler *et al*, 2018). Here, cochleae of 2-week-old WT and *Palm3*-KO littermates were chemically-cleared and immunostained with the hair cell marker Myosin7a. Then, we employed lightsheet fluorescence microscopy to enable a detailed 3D analysis of hair cell counts across the entire cochlear spiral. Myosin7a labeling served both to identify areas of interest and to establish reference points for frequency band assignment (Müller *et al*, 2005; Rankovic *et al*, 2021). Quantification of OHC survival was based on counts of Myosin7a-positive OHCs within the computed frequency bands. Compared to age-matched WT controls, *Palm3*-KO mice exhibited a significant reduction in OHC numbers across all three rows; yet, with varying degrees along the tonotopy (**Supplementary figure 5B-D**). The most extensive OHC loss occurred in the 8-12 kHz band, where survival dropped by >50% across all rows. These results indicate that loss of *Palm3* severely impacts cochlear function and OHC maintenance, warranting further investigation into the mechanisms underlying these defects.

### Genetic loss of *Palm3* results in reduced OHC length and perturbed Prestin distribution within the OHC plasma membrane

Having observed that OHCs appeared abnormally ‘compact’ in *Palm3*-KOs, we further investigated their morphology by measuring the lengths of OHCs in both pre-hearing (P11) and hearing (P21) animals to assess the developmental onset, timing and severity of the structural disturbance. Compared to WT controls, OHC length in P11 *Palm3*-KO mice was reduced by ∼22% (**Figure 3A,C**). This defect worsened in P21 *Palm3*-KO mice, with OHC length reduced by ∼40% (**Figure 3B,C’**). Even in heterozygous, *Palm3*-haploinsufficient animals (8 weeks old), we observed a ∼20% reduction in OHC length (**Supplementary figure 6**), which emphasizes the critical role of adequate Palm3 membrane levels in maintaining OHC shape.

Noting the similarities between the *Palm3* mutant phenotype – particularly the reduction in OHC length – and the known localization and knockout phenotype of Prestin (Liberman *et al*, 2002; Zheng *et al*, 2000), we conducted additional IF analyses, which confirmed the colocalization of Palm3 and Prestin within the OHC lateral plasma membrane in wildtype mice (**Supplementary figure 7**). In *Palm3*-KO OHCs, the typical homogeneous distribution of Prestin within the lateral plasma membrane was strikingly disrupted, exhibiting fragmentation, heterogeneity, and partial internalization (**Figure 3A-B**). To further assess the functional state of Prestin, we performed electrophysiological recordings on OHCs. Consistent with the observed aberrant cell lengths, we found a significant reduction in the linear capacitance (Clin) of *Palm3*-KO OHCs, indicative of a reduced membrane surface area (**Figure 3E**). Non-linear capacitance (NLC), an electrophysiological proxy of Prestin function, remained comparable to WT at P12-13, but declined at P18-20 in *Palm3*-KO OHCs (**Figure 3F**), suggesting that progressive structural deterioration leads to reduced or mislocalized Prestin. At P12-13, *Palm3-*KO OHCs displayed an elevated NLC to Clin ratio (NLC/Clin) (**Figure 3G**), indicating elevated Prestin activity per unit membrane area, likely due to normal Prestin production being confined into a progressively limited membrane space. By P18-20, NLC/Clin no longer differed from WT, indicating that Prestin packing density had reached saturation despite fragmentation. Importantly, the voltage at peak capacitance (Vh) remained consistent across all conditions, suggesting that the functional properties of Prestin were not significantly altered (**Figure 3H**). However, the slope of the bell-shaped curve (**Figure 3D,D’**) showed a significant decrease in *Palm3*-KO OHCs at P18-20, but not at P12-13 (**Figure 3I**), pointing to the possibility that Prestin activity may be slightly altered.

### Palm3 precedes Prestin arrival in the OHC lateral membrane

To investigate the developmental time course of membrane insertion for both Palm3 and Prestin, we performed IF analyses on apical turns of whole-mount organ of Corti preparations from various early postnatal stages up to the onset of hearing in mice (P0-P14, **Supplementary figure 8A**). Simultaneous co-staining for Palm3 and Prestin within the same organ of Corti was not feasible, as both antibodies were produced in rabbit. Instead, we analyzed contralateral ears of the same animals to ensure comparability of staining. In these experiments, ɑII-Spectrin was used as a common context marker in both preparations to provide a spatial reference and allow for direct comparison of Palm3 and Prestin temporal expression patterns. Palm3 labeling was prominently detectable in OHC lateral membranes as early as P0, with labeling intensity increasing by P3 and stabilizing through P7, ultimately reaching its peak intensity within the timeframe of this analysis at P14, indicating an early developmental role (**Supplementary figure 8A,C**). In contrast, Prestin labeling emerged around P3, consistent with previous studies (Oliver & Fakler, 1999; Legendre *et al*, 2008), and progressively intensified, eventually reaching its highest observed level at P14 – aligning with the maturation of OHC functionality. These findings reveal distinct temporal expression patterns between Palm3 and Prestin, with Palm3 appearing earlier in development and potentially contributing to membrane stabilization or organization required for adequate Prestin localization.

### Palm3 affects spectrin isoform expression in the OHC submembrane cytoskeleton

Based on previous research highlighting the role of paralemmins in membrane-cytoskeleton interactions (Kutzleb *et al*, 1998; Arstikaitis *et al*, 2008; Kutzleb *et al*, 2006; Nie *et al*, 2017; Sáinz-Jaspeado *et al*, 2021) and the evident subcellular localization of Palm3 in the plasma membrane compartment, we asked whether the recently described interaction of Palm1 with βII-Spectrin in the periodic submembrane cytoskeleton of neuronal axons (Macarrón-Palacios *et al*, 2025) could – by analogy – reflect a conserved paralemmin/spectrin association across isoforms. First, we identified the most likely candidate spectrin isoforms by utilizing the publicly available transcriptome profiling data for OHCs from the gEAR database (**Supplementary figure 9**; Orvis *et al*, 2021; https://umgear.org/), revealing αII-, βI- and βII-Spectrin, but not αI-, βIV- and βV-Spectrin to be detected in hair cells. These three isoforms were therefore selected for further validation by immunohistochemistry and confocal microscopy. The co-labelings with Palm3 confirmed that αII-, βI- and βII-Spectrin indeed co-localize with Palm3 at the lateral walls of auditory hair cells (**Figure 4A-C’**). Additionally, all three spectrin isoforms are found to be enriched in the actin-rich cuticular plates. Strikingly βI-Spectrin – a major component of the erythrocytic membrane cytoskeleton – was robustly localized to the lateral plasma membranes of IHCs and OHCs (**Figure 4B-B’**), with erythrocytes in blood vessels of the organ of Corti serving as an internal positive control for target specificity (**Supplementary figure 10**).

**Figure 4.**
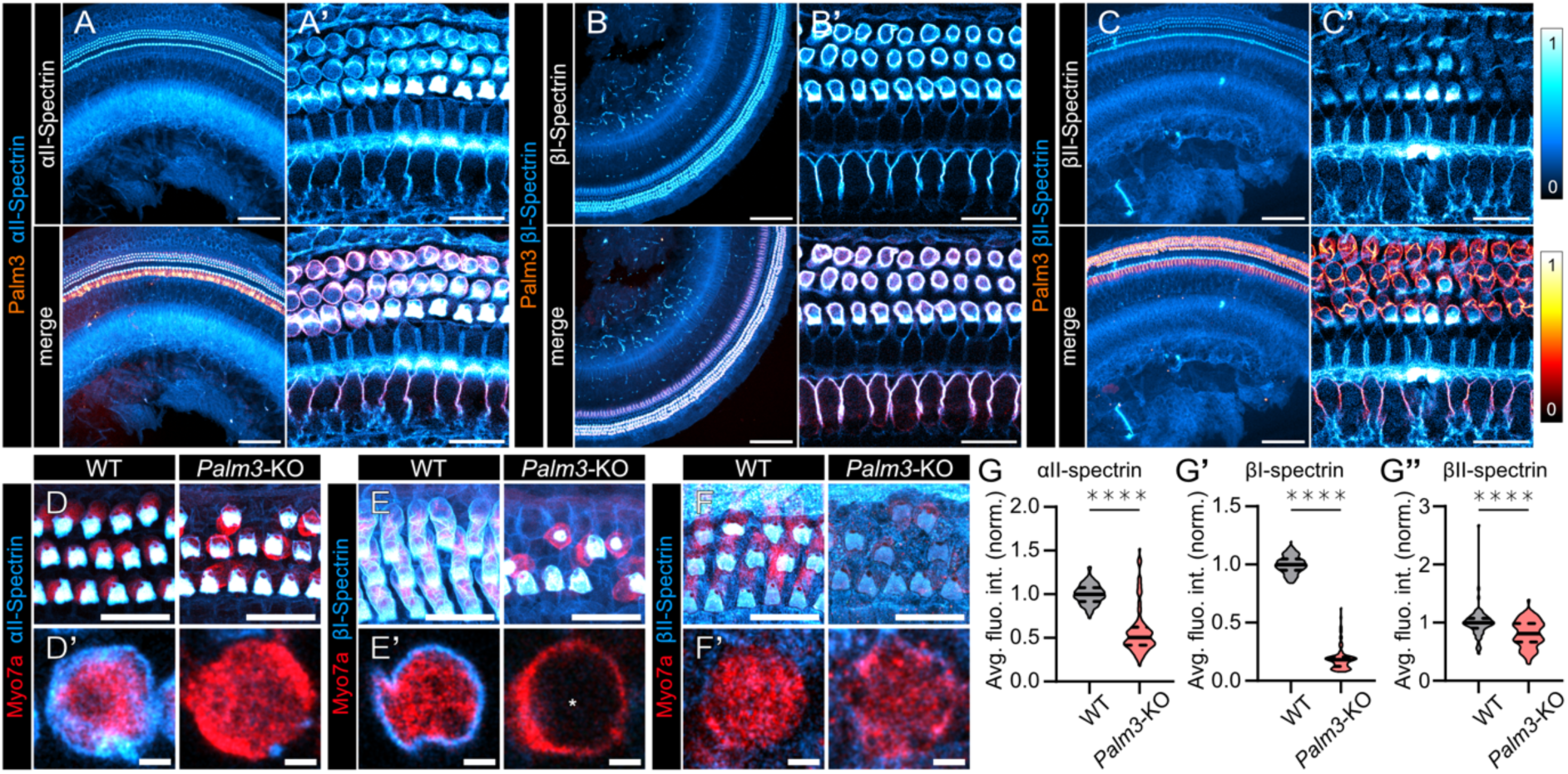
Loss of Palm3 reduces spectrin abundance at OHC lateral plasma membranes. (A-C’) Representative confocal maximum z-projections showing localization of Palm3 (red hot) and either αII-Spectrin, βI-Spectrin or βII-Spectrin (cyan hot) within the organ of Corti. (A,B,C) Overview images are depicted on the left columns (scale bars: 100 µm), whereas (A’,B’,C’) right columns show single optical sections of the respective staining at higher magnification (scale bars: 20 µm). (D-F’) Representative confocal maximum z-projections of OHCs stained against Myosin7a (red) and either αII-Spectrin, βI-Spectrin or βII-Spectrin (cyan hot) from 6-week-old WT and *Palm3*-KO. (D,E,F) Upper panels show overview of the OHC rows and (D’,E’,F’) lower panels show single optical sections of individual OHCs of the respective stacks. Scale bars: upper panels: 20 µm, lower panels: 2 µm. The * in E’ indicates the unstained nucleus. (G-G”) Violin plots showing average fluorescence intensity of (G) αII-Spectrin (*N*_animals_= 5, *n*_Corti_ = 10, *n*_OHC_ = 134 for WT and *N*_animals_= 5, *n*_Corti_ = 10, *n*_OHC_ = 90 for *Palm3*-KO), (G’) βI-Spectrin (*N*_animals_= 5, *n*_Corti_ = 10, *n*_OHC_ = 130 for WT and *N*_animals_= 5, *n*_Corti_ = 10, *n*_OHC_ = 88 for *Palm3*-KO) and (G”) βII-Spectrin (*N*_animals_= 3, *n*_Corti_ = 6, *n*_OHC_ = 71 for WT and *N*_animals_= 3, *n*_Corti_ = 6, *n*_OHC_ = 68 for *Palm3*-KO) along the OHC lateral wall is decreased in absence of Palm3. Data information: Two-tailed Mann-Whitney test: **** *P* < 0.0001.

We next assessed whether Palm3 deficiency alters the abundance of individual spectrins by measuring fluorescence intensities in the lateral wall of *Palm3*-KO OHCs. We found that the average fluorescence intensity of αII-Spectrin was reduced by ∼42% in *Palm3*-KO OHC lateral membranes (**Figure 4G**), while βI-Spectrin exhibited a dramatic ∼80% decrease, indicating a strong Palm3 dependency for its stability and/or correct localization (**Figure 4G’**). Lastly, in line with previous work (Liu *et al*, 2019) βII-Spectrin, which generally showed the weakest signal intensity in the lateral membrane but is enriched in the cuticular plates, declined by ∼20% in fluorescence intensity (**Figure 4G”**). In contrast to our observations at the lateral membranes, average fluorescence intensities in the cuticular plates showed largely opposing trends: while αII-Spectrin and βI-Spectrin intensity increased by ∼49% and ∼15% in absence of Palm3 respectively, βII-Spectrin declined by ∼16% (**Supplementary figure 11C**). These reciprocal shifts imply that *Palm3* deletion differentially affects the targeting to or stabilization within the submembraneous cytoskeleton, thus leading to compensatory re-distribution of spectrin isoforms between different compartments.

To define the relationship between Palm3 and these three spectrin isoforms within the organ of Corti at higher spatial resolution, we used two-color 2D stimulated emission-depletion (2D-STED) optical nanoscopy on whole-mount preparations of the organ of Corti. In these experiments, samples were co-labeled with antibodies against Palm3 and each of the spectrin isoforms – αII-, βI-, or βII-Spectrin – then embedded in melamine and sectioned into ∼80-90 nm ultrathin slices (Revelo *et al*, 2014). This approach physically minimizes the third dimension of the sample, and hence allowed us to capitalize on the maximum lateral resolution of 2D-STED that would not be achievable in intact preparations due to tissue-based spherical aberrations. This revealed unique staining patterns for Palm3 and the various spectrin isoforms: Palm3 signal showed a striking punctate distribution along the OHC membrane (**Figure 5A’-C’**), while αII-Spectrin and βI-Spectrin exhibited similar, yet somewhat more continuous, staining patterns along the OHC membrane (**Figure 5A’,B’**). In contrast, βII-Spectrin displayed a broader labelling distribution, detected in both the OHC plasma membrane and cytosol, but was mainly concentrated within the cuticular plates (**Figure 5C,C’**). Interestingly, while all spectrin staining patterns closely aligned with Palm3 along the longitudinal as well as circumferential plasma membrane, βI-Spectrin in particular appeared to closely follow the longitudinal staining pattern of Palm3 (**Figure 5A”’-C”’**). Notably, compared to Palm3 IF, the spectrin IF distribution appeared slightly shifted towards the intracellular space (**Figure 5A”-C”**).

**Figure 5.**
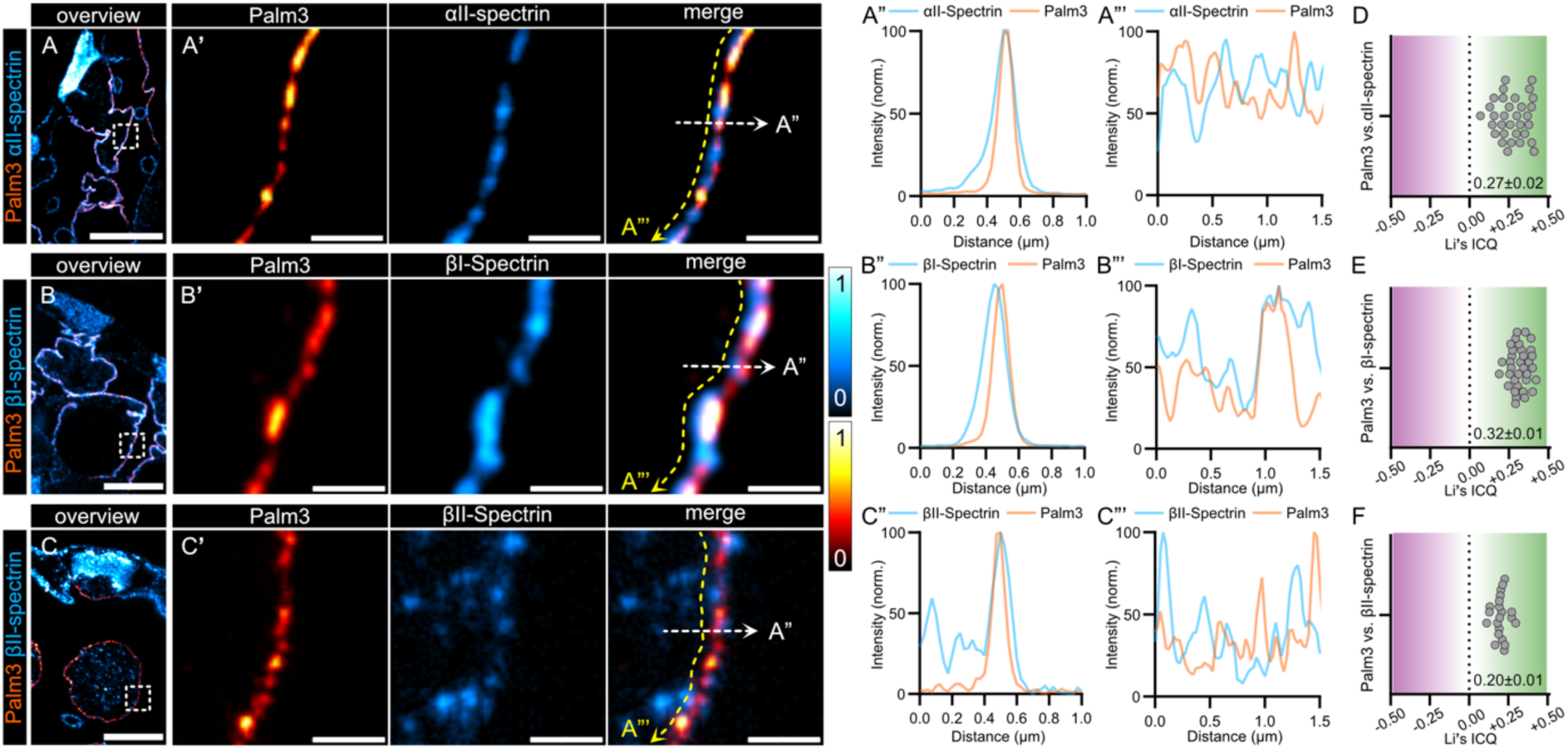
Localization of Palm3 and different spectrin isoforms in the OHC lateral plasma membrane. (A-C) Two-color 2D-STED IF images of ∼80-90 nm ultrathin melamine-embedded sections of organs of Corti from 6-8-week-old WT mice, showing a single OHC labeled with Palm3 (red hot) and either (A) αII-Spectrin, (B) βI-Spectrin, or (C) βII-Spectrin (cyan hot). Scale bars: 5 µm. (A’-C’) Close-up views of insets depicted in (A-C), showing a region of interest in the OHC lateral plasma membrane labeled with Palm3 (red hot) and either (A’) αII-Spectrin, (B’) βI-Spectrin, or (C’) βII-Spectrin (cyan hot). Dotted line arrows indicate the starting and end point of the intensity profile analyses shown in (A”-C” and A”’-C”’). Scale bars: 0.5 µm. (A”-C”) Transverse intensity profiles of Palm3 (orange) and either (A”) αII-Spectrin, (B”) βI-Spectrin, or (C”) βII-Spectrin (blue). (A’”-C’”) Longitudinal intensity profiles of Palm3 (orange) and either (A’”) αII-Spectrin, (B’”) βI-Spectrin, or (C’”) βII-Spectrin (blue). (D-F) Scattered dot plots showing Li’s intensity correlation quotient (ICQ) values highlights the spatial correlation between Palm3 and either (D) αII-Spectrin (*N*_animals_= 3, *n*_images_= 32), (E) βI-Spectrin (*N*_animals_= 3, *n*_images_= 34), or (F) βII-Spectrin (*N*_animals_= 2, *n*_images_= 21). Mean ± SEM values are depicted in the Figure.

To quantify the spatial relationship between Palm3 and each respective spectrin isoform, we calculated Li’s intensity correlation quotient (ICQ), which tests the correlation between individual pixel intensities of two distinct staining patterns (Li *et al*, 2004). The resulting ICQ value falls within a range of −0.5 and +0.5, where positive values indicate the degree of colocalization or dependency, values near zero suggest random distribution, and negative values imply segregation or mutual exclusion. ICQ analyses revealed positive ICQ values for Palm3 and all tested spectrins, indicating a significant degree of staining dependency between these proteins – in particular with αII- and βI-Spectrin (**Figure 5D-F**).

### Palm3 appears periodically spaced in the OHC plasma membrane

Next, to obtain a better understanding of the Palm3 cluster distribution within the nano-architecture of the OHC lateral membrane, we analyzed the nearest-neighbor distances (NND) of Palm3 immunofluorescence puncta in our 2D-STED images. In longitudinally-sectioned OHCs, the average Palm3 NND was 143 ± 1.5 nm, whereas in cross-sectioned OHCs, representing circumferential distribution, the average NND was slightly shorter at 132 ± 1.5 nm (*N*animals= 4; *n*images= 6; *n*Palm3= 953 for longitudinal; *N*animals= 4; *n*images= 8; *n*Palm3= 891 for cross section). In a few cells, where the OHC plasma membrane was sectioned slightly angled to enable an *en face* view (**Figure 6A,A’**), we could extend this analysis to measure the NNDs of up to five of the closest surrounding Palm3 clusters (1^st^ NND = 124 ± 0.8 nm, 3^rd^ NND = 161 ± 0.9 nm, 5^th^ NND = 194 ± 1.1 nm; **Figure 6B**). Similarly, we could perform the same type of analysis with αII-Spectrin (1^st^ NND = 138 ± 2.4 nm, 3^rd^ NND = 188 ± 2.4 nm, 5^th^ NND = 229 ± 2.6 nm; **Figure 6C**) and found that the observed distances are well within the range of β-Spectrin spacing in neuronal axons – where spectrin IF exhibits a characteristic periodicity pattern at ∽180-190 nm (Pan *et al*, 2018; Han *et al*, 2017; D’Este *et al*, 2015). Collectively, these findings strongly indicate that (i) Palm3 co-localizes with the submembraneous actin/spectrin cytoskeleton, where it – based on ICQ analysis – (ii) exerts interdependency in particular with αII- and βI-Spectrins and (iii) appears to be evenly spaced along the OHC circumference – albeit with a mild longitudinal stretch. Hence, these findings further support the hypothesized molecular association between Palm3 and spectrin isoforms.

**Figure 6.**
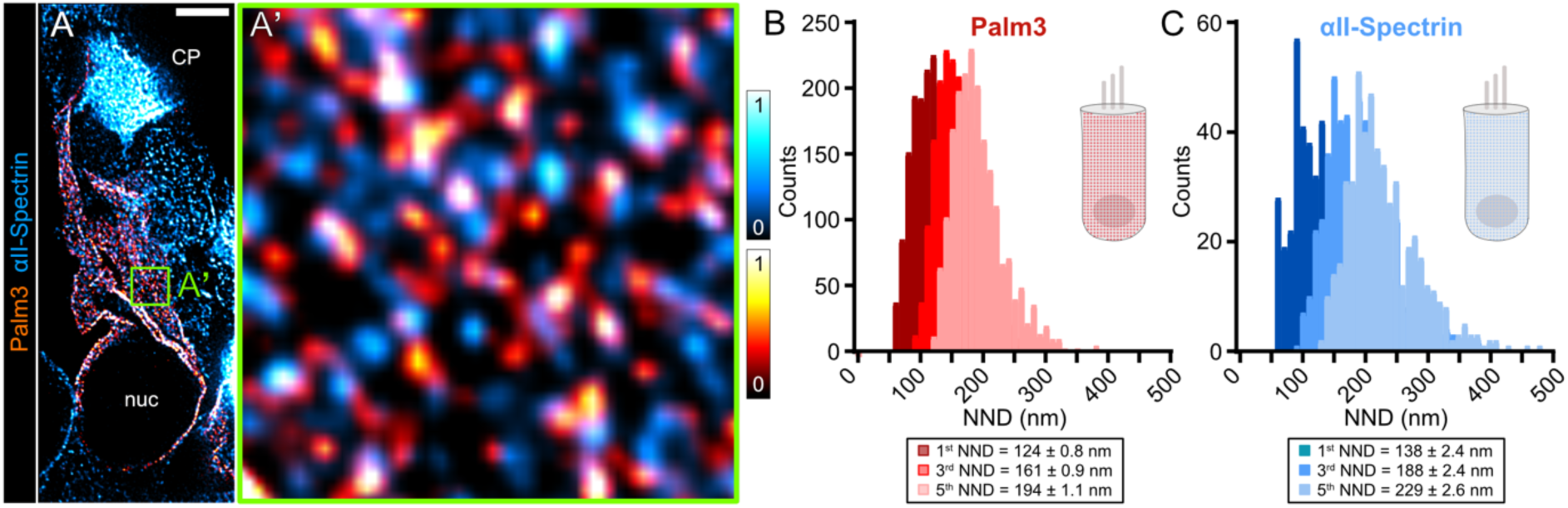
Spatial distribution of Palm3 and αII-Spectrin within the OHC lateral plasma membrane. (A,A’) Representative two-color 2D-STED IF images of ∼80-90 nm ultrathin melamine-embedded sections of Palm3-stained organs of Corti from 6-8-week-old WT mice, showing (A) a single OHC and (A’) a close up view of the inset depicted in (A) labeled with Palm3 (red hot) and αII-Spectrin (cyan hot). Scale bar in (A) 2 µm, box width in (A’) 1.5×1.5 µm. (B) First to fifth NNDs between Palm3 immunostaining puncta. Inset shows the mean NND value for each configuration. *N*_animals_= 4; *n*_images_= 9; for 1^st^ NND: *n*_Palm_= 1965, for 3^rd^ NND: *n*_Palm_= 1966, and for 5^th^ NND: *n*_Palm_= 1966. (C) First to fifth NNDs between αII-Spectrin immunostaining puncta. Inset shows the mean NND value for each configuration. *N*_animals_= 4; *n*_images_= 9; for 1^st^, 3^rd^, 5^th^ NND: *n*_αII-Spectrin_= 504.

### Loss of Palm3 disrupts the structure of the subsurface cisternae underneath the cortical lattice

To examine potential ultrastructural alterations in the lateral membrane of *Palm3*-KO OHCs, we next employed electron tomography on high-pressure frozen, freeze-substituted (HPF/FS) apical turns of the organ of Corti (Chakrabarti *et al*, 2018, 2022). This method enables high-resolution imaging while preserving near-native structural integrity, rendering it well-suited to compare the lateral wall ultrastructure of WT and *Palm3*-KO OHCs (**Figure 7A-B”**). Indeed, the tomographic reconstructions of the supranuclear SSCs and proximal plasma membrane revealed striking differences between genotypes: apical OHCs in *Palm3*-KO mice exhibited a substantial decrease in supranuclear SSC abundance along the lateral wall (**Figure 7C**). The few residual SSCs in *Palm3*-KO OHCs were positioned more distant from the plasma membrane compared to those in WT OHCs (**Figure 7D**), suggesting disrupted spatial arrangement – potentially due to impaired SSC anchoring within the lateral wall complex. Morphometric analysis also showed that SSCs in *Palm3*-KO OHCs appeared more flattened and structurally simplified, as indicated by reduced surface-to-volume ratios (**Figure 7E**). Beyond the SSC, mitochondria – which typically located proximal to the plasma membrane – were found at greater distances in *Palm3*-KO OHCs. This was confirmed by quantitative analysis, which showed a significant increase in the average distance between mitochondria and the plasma membrane (**Figure 7F**).

**Figure 7.**
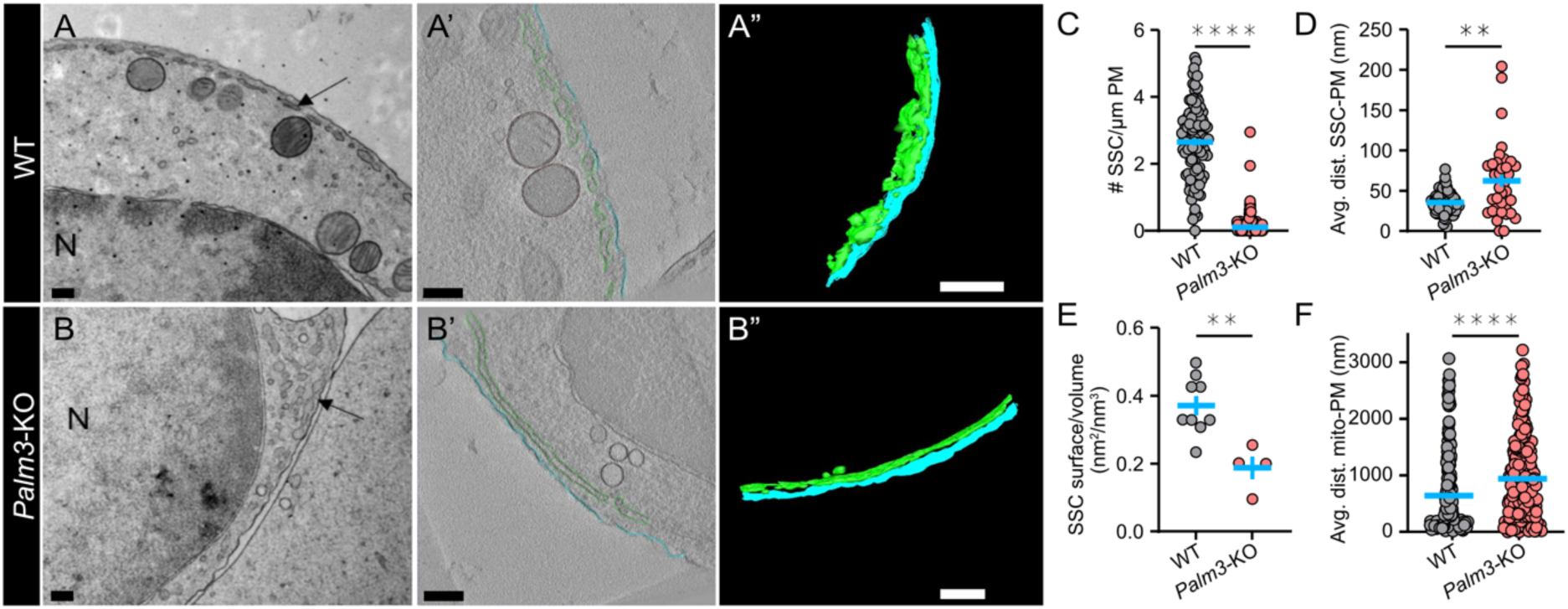
Palm3-KO subsurface cisternae (SSC) are fewer in number, structurally less complex and located more distally from the lateral plasma membrane. (A-B) Representative transmission electron microscopy (TEM) micrograph of (A) WT and (B) *Palm3*-KO OHC lateral wall obtained after high pressure freezing/freeze-substitution from P14 mice. N indicates the cell nucleus and arrow indicates SSCs. Scale bars: 0.2 µm. (A’-B”) Representative virtual electron tomographic sections with their (A”,B”) corresponding 3D models of (A’,A”) WT and (B’,B”) *Palm3*-KO OHC lateral wall obtained after high pressure freezing/freeze-substitution from P14 mice. The tomographic reconstructions show the OHC plasma membrane in cyan and SSC in green. Scale bars: 0.2 µm. (C-E) Scattered dot plots display the quantification of (C) number of SSC per µm membrane (*N*_animals_= 2, *n*_OHCs_ = 23, *n*_measurements_= 102 for WT and *N*_animals_= 5, *n*_OHCs_ = 30, *n*_measurements_= 165 for *Palm3*-KO), (D) average distance of SSC to the plasma membrane (PM) (*N*_animals_= 2, *n*_OHCs_ = 23, *n*_measurements_= 81 for WT and *N*_animals_= 2, *n*_OHCs_ = 21, *n*_measurements_= 36 for *Palm3*-KO), and (E) surface per volume mesh membrane (*N*_animals_= 2, *n*_OHCs_ = 23, *n*_measurements_= 9 for WT and *N*_animals_= 2, *n*_OHCs_ = 21, *n*_measurements_= 4 for *Palm3*-KO). Blue bars represent mean ± SEM. (F) Average distance of mitochondria to the plasma membrane is increased in *Palm3*-KO OHCs (*N*_animals_= 2, *n*_OHCs_ = 23, *n*_measurements_= 230 for WT and *N*_animals_= 2, *n*_OHCs_ = 21, *n*_measurements_= 255 for *Palm3*-KO). Data information: Two-tailed Mann-Whitney test: ** *P* < 0.01, **** *P* < 0.0001; Mean ± SEM.

### AAV-mediated *Palm3*-rescue partially restores hearing loss in *Palm3*-KO mice

Given that hair cell degeneration in *Palm3*-KO mice begins around the onset of hearing and progresses over time, we evaluated whether early postnatal viral rescue could prevent the pathophysiological consequences of *Palm3* loss. We used an adeno-associated viral vector (AAV.S; Ivanchenko *et al*, 2021) carrying a YFP-tagged *Palm3* fusion gene under the control of the human *Myo15a* promoter (*AAV-S_Myo15a_YFP-Palm3*; **Figure 8A**; **Supplementary figure 12**). The vector was delivered via unilateral intracochlear injection through the round window membrane of *Palm3*-KO mouse pups at P4-6 (Rankovic *et al*, 2021). This method enabled within-subject comparisons between injected (ipsilateral) and non-injected (contralateral) ears to assess the functional and morphological magnitude of rescue efficacy. Auditory function and cochlear morphology were then evaluated 4-5 weeks post-injection (**Figure 8B**): *Palm3*-rescue mice exhibited modest hearing improvements (**Figure 8C-E**). Specifically, ABR amplitudes in response to 100 dB and 20 Hz click stimulation as well as pure-tone thresholds were mildly but statistically significantly improved in the injected ears (**Figure 8C-D**). DPOAEs at the f2 frequency of 11.3 kHz show a trend towards increased amplitudes in the injected ears, although this effect did not reach statistical significance (**Figure 8E**).

**Figure 8.**
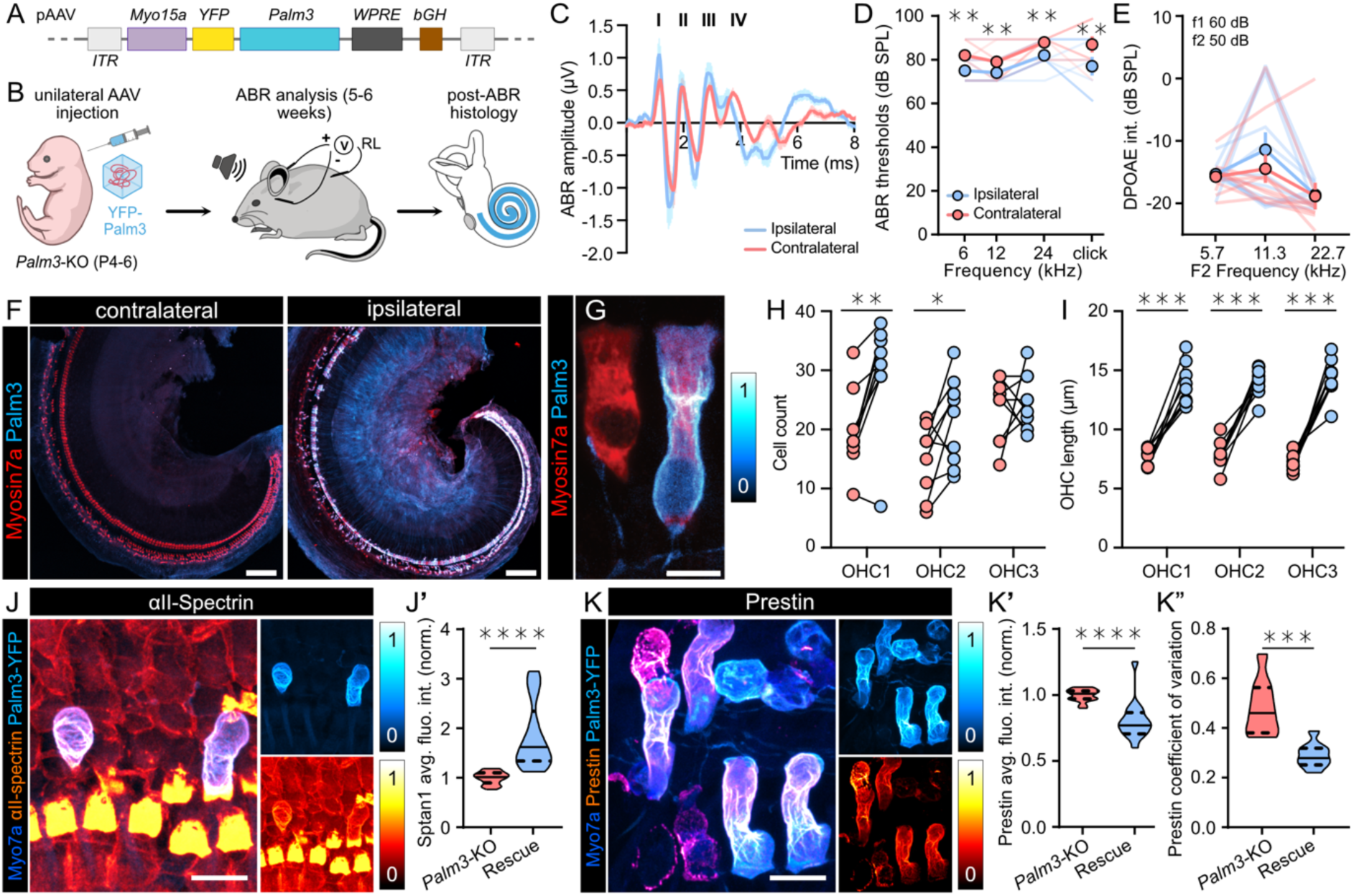
AAV-mediated rescue improved hearing function and restored OHC counts and lengths in Palm3-KO mice. (A) The vector is based on a pAAV backbone and carries the mouse *Palm3* gene fused to a yellow fluorescent protein gene (*YFP*). This fusion construct is inserted between the human *Myo15a* promoter and a non-oncogenic variant of Woodchuck hepatitis virus posttranslational regulatory element (*WPRE*), followed by the bovine growth hormone (*bGH*) polyadenylation signal sequences. The entire cassette is flanked by inverted terminal repeats (*ITR*s). (B) Schematic overview of the AAV-mediated Palm3 rescue experimental procedure. *AAV.S_Myo15a_YFP-Palm3* was administered via round window intracochlear injection at P4-6 in Palm3-KO mouse pups. After the injection, ABR and DPOAE measurements were performed to evaluate hearing function following a maturation period of 4 to 5 weeks. Subsequently, the animals were sacrificed, and their cochleae harvested for morphological assessment. (C) Grand averages of ABR waveforms upon 100 dB and 20 Hz click stimulation in 5-week-old *Palm3-*KO mice following injection of *AAV.S_Myo15a_YFP-Palm3* in ipsilateral ears (blue) and non-injected contralateral ears (pink) of the same animals. Averages represented by lines, means ± SEM represented by shaded areas, ABR waves generated along the auditory pathway represented by roman numerals (Jewett Waves I-V). (D) ABR thresholds to clicks and 6, 12, and 24 kHz tone bursts in injected and contralateral ears (*N*_animals_= 10, with *n*_Corti_ = 10 for ipsilateral and *n*_Corti_ = 10 for contralateral; Two-way ANOVA with Holm-Šídák‘s multiple comparisons correction: ** *P* < 0.01; Mean ± SEM). (E) Slight increase in DPOAE amplitude at f2 frequency of 11.3 kHz suggests a subtle improvement of amplification in *Palm3*-rescue. Orange and light blue lines indicate the average noise floor (*N*_animals_= 10, with *n*_Corti_ = 10 for ipsilateral and *n*_Corti_ = 10 for contralateral; Two-way ANOVA with Holm-Šídák‘s multiple comparisons correction: *P* = 0.6225; Mean ± SEM). (F) Representative confocal maximum z-projections of whole-mount apical turns of non-injected (contralateral-) and injected (ipsilateral) ears of 5-week-old *Palm3*-KO animals. Cortis were stained for Myosin7a (red) and Palm3-YFP (cyan hot). Scale bars: 100 µm. (G) A *Palm3*-transduced OHC displays improved cellular morphology in comparison to a neighboring non-transduced OHC. Cortis stained for Myosin7a (red) and Palm3-YFP (cyan hot). Scale bar: 10 µm. (H) Single value plots showing OHCs counts within a 300 µm x 300 µm region of interest to exhibit a trend toward increased survival in *Palm3-*rescue ears (*N*_animals_= 9, with *n*_Corti_ = 9 for ipsilateral and *n*_Corti_ = 9 for contralateral: Two-way ANOVA with Holm-Šídák‘s multiple comparisons correction: ns = not significant; * *P* < 0.05; ** *P* < 0.01; Mean ± SEM; connecting lines identify individual animals). (I) *Palm3-*transduced OHCs show restored cell lengths in all three OHC rows (*N*_animals_= 9, with *n*_Corti_ = 9 for ipsilateral and *n*_Corti_ = 9 for contralateral; Two-way ANOVA with Holm-Šídák‘s multiple comparisons correction: **** *P* < 0.0001; Mean ± SEM). (J) Representative confocal maximum z-projections of OHCs in AAV-injected, 5-week-old *Palm3*-KO animals (n = 5). OHCs were stained with Myosin7a (blue), Palm3 (cyan hot), and αII-Spectrin (red hot). Scale bars: 5 µm. (J’) Violin plot showing normalized average fluorescence intensity of αII-Spectrin revealed restored αII-Spectrin-labeling intensity in *Palm3*-transduced OHCs (*N*_animals_= 5, with *n*_OHCs_ = 27 for Palm3-negative and *n*_OHCs_ = 25 for Palm3-positive; Two-tailed Mann-Whitney test: **** P < 0.0001; Mean ± SEM). (K) OHCs were stained with Myosin7a (blue), Palm3 (cyan hot), and Prestin (red hot). Representative confocal maximum z-projections of OHCs in AAV-injected, 5-week-old *Palm3*-KO animals (n = 5). Scale bars: 5 µm. (K’) Violin plot showing normalized average fluorescence intensity of Prestin revealed decreased Prestin-labeling intensity in *Palm3*-transduced OHCs. (*N*_animals_= 4, with *n*_OHCs_ = 17 for Palm3-negative and *n*_OHCs_ = 15 for Palm3-positive; Two-tailed Mann-Whitney test: **** P < 0.0001; Mean ± SEM). (K”) Coefficient of variation of Prestin-labeling intensity indicates a restoration of homogeneous Prestin distribution over the lateral membrane of OHCs in *Palm3*-transduced OHCs. (*N*_animals_= 4, with *n*_OHCs_ = 17 for Palm3-negative and *n*_OHCs_ = 15 for Palm3-positive; Two-tailed Mann-Whitney test: *** P < 0.001; Mean ± SEM).

### OHC structure and survival rate are partially restored in AAV-rescued *Palm3*-KO mice

To evaluate the structural and cellular effects of AAV-mediated *Palm3* rescue, cochleae were collected post-ABR recordings and processed for subsequent IF analysis (**Figure 8F**). We observed notable variability in OHC transduction efficiency, while IHCs consistently exhibited higher transduction and expression rates (**Supplementary figure 13A**). This heterogeneity – together with lack of target cell specificity – is probably responsible for the modest rescue success at the systems level (ABR).

To determine whether viral *Palm3* re-introduction enhanced hair cell survival in *Palm3-*KO mice, we quantified hair cells within a defined 300 x 300 µm region of the apical cochlear turn of the organ of Corti in *Palm3*-rescue animals. IHC counts showed a slight reduction of ∼11.6% in the ipsilateral ear (**Supplementary figure 13B**) which is more pronounced than the weak impact of the *Palm3*-KO itself at 7-8 weeks (∼6%, WT: 25.0 ± 1.16; *Palm3*-KO: 23.5 ± 0.22; **Figure 2B**) – presumably due to cytotoxic effects from aberrant Palm3 expression or to viral (over-) load, which appear to disrupt IHC morphology and compromise cellular structural homeostasis (**Supplementary figure 13C**). In contrast, in OHCs which are severely affected by the *Palm3*-KO, a beneficial effect of the rescue prevailed: OHC survival in the ipsilateral ears trended upwards, particularly in the innermost (OHC1, ∼53%) and middle rows (OHC2, ∼41%) (**Figure 8H**).

To assess whether the structural integrity of OHCs was restored in *Palm3*-transduced cells, we further examined cellular length. Transduced OHCs displayed elongated and creased plasma membranes, resembling the wildtype morphology (**Figure 8G**). Quantitative assessment of OHC length confirmed length restoration in ipsilateral ears, reaching values comparable to OHCs in P21 WT mice (**Figure 8I**; **Figure 3C’**). Together, these results suggest that AAV-mediated *Palm3* delivery in *Palm3*-KO mice generally supports improved OHC survival and facilitates structural restoration, thus resulting in the functional improvements observed in **Figure 8C-E**.

### Viral Palm3 rescue reestablishes αII-Spectrin and Prestin abundance in *Palm3*-KO OHCs

To further assess the cellular effects of *Palm3* rescue on the plasma membrane scaffold of *Palm3*-KO OHCs, we examined whether the expression of αII-Spectrin could be restored. As αII-Spectrin heterodimerizes with βI-as well as βII-Spectrins, it served as a surrogate to assess spectrin scaffold integrity. Quantitative IF analysis revealed a statistically significant increase in αII-Spectrin average fluorescence intensity in *Palm3*-transduced OHCs compared to neighboring non-transduced OHCs within the same organ of Corti (**Figure 8J,J’**). These results suggest that *Palm3* re-introduction restores αII-Spectrin expression in the lateral membrane.

Finally, we investigated whether *Palm3* rescue would also restore the irregular and clustered distribution of Prestin in *Palm3*-KO OHCs (**Figure 3B**). Indeed, Prestin staining appeared visually more homogeneous relative to that in non-transduced OHCs (**Figure 8K**). Quantitative analysis revealed a slight decrease in average Prestin fluorescence intensity in *Palm3*-transduced OHCs, likely due to the loss of abnormally high-intensity IF signal clusters and a transition toward a more homogeneous distribution as observed in wildtype OHCs (**Figure 8K’**). Supporting this, the coefficient of variation (CV) of Prestin IF puncta intensity was significantly higher in *Palm3-*KO than in *Palm3*-recued OHCs, indicating greater spatial heterogeneity in Prestin distribution in the KOs (**Figure 8K”**). Conversely, the lower CV observed in *Palm3*-transduced OHCs suggests a restoration of a more uniform Prestin distribution along the lateral membrane.

## Discussion

In the present study, we investigated the functional and morphological consequences of *Palm3* deficiency within the peripheral auditory system. Alongside systems-physiological analyses, we employed immunohistochemistry in combination with a range of advanced microscopy techniques – including lightsheet fluorescence microscopy, confocal and 2D-STED microscopy, as well as electron tomography – to elucidate the consequences of *Palm3* loss on the morphology and function of OHCs. Our findings reveal that *Palm3*-deficient mice exhibit early-onset, progressive hearing loss that was associated with gradual degeneration of auditory hair cells along the entire tonotopic axis of the cochlea. Palm3 localizes specifically to the lateral plasma membranes of IHCs and OHCs, where it stabilizes the cortical lattice. In OHCs, in particular, the absence of *Palm3* leads to cell length reduction, disrupted distribution of Prestin, misdistribution of spectrins, and a decrease in the number and structural integrity of subsurface cisternae (SSCs) along the OHC lateral plasma membrane. AAV-mediated rescue of *Palm3*-KO animals partially restored auditory function, due to enhanced OHC survival and restoration of OHC lengths, and re-establishment of ɑII-Spectrin expression and proper Prestin localization. Together, these findings highlight a critical role for Palm3 in maintaining the structural integrity and function of auditory hair cells and demonstrate its potential as a novel deafness gene.

### Genetic loss of *Palm3* disrupts cochlear amplification and OHC integrity by compromising cytoskeletal and membrane stability

Genetic loss of Palm3 leads to a substantial ∼50 dB SPL elevation in auditory thresholds and a near-complete loss of active cochlear amplification, indicating its critical role in peripheral auditory processing. Localized to the lateral plasma membrane of auditory hair cells, Palm3 is essential for maintaining hair cell integrity – particularly in OHCs. This heightened dependency of OHCs on Palm3 is likely due to their specialized role in voltage-driven somatic electromotility and the concomitant demand on the stability of their lateral cell wall (Borkó *et al*, 2005; Batta *et al*, 2003; Hallworth, 1995; He & Dallos, 2000).

The severe vulnerability of *Palm3*-KO OHCs is reminiscent of the phenotype observed in *Palmd*-KO endothelial cells, where reduced mechanical resilience via disrupted nuclear actin caps contributes to pathology (Sáinz-Jaspeado *et al*, 2021). Similarly, OHCs highly depend on a well-organized actin/spectrin cytoskeleton to support both their somatic electromotility and structural integrity (Brownell, 1990; Holley *et al*, 1992; Oghalai *et al*, 1998; Holley & Ashmore, 1990; Wada *et al*, 2004). In *Palm3*-KO OHCs, mis-distribution of ɑII-, βI- and βII-Spectrins from the lateral plasma membrane to the cuticular plate indicates destabilization of the submembrane skeleton (Yao *et al*, 2022; Liu *et al*, 2019), while abnormalities in SSCs and misplacement of membrane-proximal mitochondria indicate that the disruption of the submembrane skeleton also perturbs the ultrastructural organization of these endomembrane compartments (Lue & Brownell, 1999; Perkins *et al*, 2020) and possibly OHC energetics (Song & Santos-Sacchi, 2015). While Prestin distribution is altered in *Palm3*-KO OHCs, their NLC remains largely preserved in young pre-hearing mice, suggesting that electromotility *per se* is not affected. Therefore, degeneration likely arises from compromised mechanical resilience due to aberrant cytoskeletal and ultrastructural integrity that renders OHCs vulnerable to recurrent membrane distortion. In contrast, IHCs – which do not engage in somatic motility – exhibit milder structural defects, indicating a lesser reliance on these same membrane characteristics (Ashmore, 2008; Peng & Ricci, 2011). Future studies – e.g., involving chronic sound deprivation or excessive noise during rearing – could clarify whether mechanical stress is the principal driver of OHC degeneration.

Building on the observed cytoskeletal and lateral membrane disorganization in *Palm3*-KO OHCs, we examined whether such structural abnormalities might also impact the localization and functional stability of Prestin over time. The striking overlap in localization patterns between Palm3 and Prestin, along with the similar reduction in OHC lengths observed in both *Palm3*- and *Prestin*-KO mice, suggest a functional connection between the two proteins. In *Palm3*-KO mice, Prestin becomes aberrantly distributed within the OHC lateral membrane. Despite this mislocalization, NLC remains unaltered in younger animals, indicating preserved voltage-dependent behavior and blanket distribution of Prestin at early developmental stages. In contrast, the significant reduction in NLC observed in older *Palm3*-KO mice suggests a progressive decline in electromotility. Such a decline has been previously linked to Prestin-associated factors including direct functional impairment (Liberman *et al*, 2002; He *et al*, 2009; Dallos *et al*, 2008), altered protein expression (Cheatham *et al*, 2015; Seymour *et al*, 2016; Abe *et al*, 2007), or disrupted membrane localization (Zheng *et al*, 2005; Sturm *et al*, 2007). In our study, the fragmented and patchy distribution of Prestin within the lateral membrane of *Palm3*-KO OHCs, which became increasingly pronounced with advancing age, likely contributes to the observed disruption of NLC in the older animals.

Further analysis of NLC/Clin ratio revealed elevated values in younger *Palm3*-KO OHCs compared to WT, suggesting relatively higher Prestin activity per unit membrane area. This likely reflects normal Prestin production while the membrane surface area decreases, leading to greater protein packing density within the available membrane space. In older *Palm3*-KO mice, the ratio resembled that of wildtype values, suggesting that Prestin had reached its maximal packing density. Yet, despite the uneven membrane distribution of Prestin observed in *Palm3*-KO OHCs, the overall Prestin levels appear comparable to those in WT OHCs. Supporting this, Vh – a reliable indicator of Prestin functional maturation (Oliver & Fakler, 1999; Santos-Sacchi & Tan, 2018) – remained stable across age groups and genotypes, indicating that Prestin protein function as such is largely preserved in *Palm3*-KO OHCs. Taken together, our findings indicate that while Palm3 is not essential for the core function or expression of Prestin, it plays a critical role in maintaining proper membrane organization and distribution of Prestin over time.

### Palm3 contributes to the stability and maintenance of the OHC submembrane cytoskeleton

Consistent with earlier work implicating other Paralemmin isoforms in membrane-cytoskeleton dynamics (Kutzleb *et al*, 1998; Arstikaitis *et al*, 2008; Sáinz-Jaspeado *et al*, 2021; Macarrón-Palacios *et al*, 2025), we observed that *Palm3* deletion leads to a significant reduction in the lateral membrane localization of ɑII-, βI- and βII-Spectrins. These findings underscore the role of Palm3 in maintaining submembrane cytoskeleton integrity within this specialized plasma membrane domain. The varying degrees of signal loss across the investigated spectrin isoforms further suggest differential regulatory mechanisms and potentially isoform-specific interactions or dependencies for membrane recruitment and/or stabilization. Interestingly, we found the effects of Palm3 loss to extend beyond the lateral wall. The reduced βII-Spectrin intensity in the cuticular plate of *Palm3*-KO OHCs raises the possibility that Palm3 may contribute to spectrin organization across distinct cytoskeletal compartments, potentially through indirect mechanisms such as trafficking or scaffolding. In contrast, the observed increase of ɑII- and βI-Spectrin in the cuticular plates of *Palm3*-KO OHCs may reflect a compensatory response to impaired membrane anchoring within the lateral wall. Whether this effect involves direct interaction with spectrins or is mediated through intermediary scaffolding proteins remains to be clarified.

Using 2D-STED microscopy on ultrathin melamine-embedded sections, we observed that Palm3 exhibits a membrane-associated spacing of ∼124-194 nm in OHCs, with distances slightly varying between longitudinal and circumferential orientations. This pattern falls within the range reported for β-Spectrin periodicity and length in neurons and erythrocytes (Pan *et al*, 2018; Zhong *et al*, 2014; Bennett *et al*, 1982; Shotton *et al*, 1979; D’Este *et al*, 2015, 2017), raising the possibility that Palm3 aligns with spectrins to orchestrate the membrane-cytoskeleton architecture. However, unlike the well-organized linear arrays seen in axons (Macarrón-Palacios *et al*, 2025; D’Este *et al*, 2015), the OHC lateral wall is subject to dynamic mechanical changes during electromotility, likely resulting in a more complex and flexible cytoskeletal meshwork (Holley, 1996). This distinction may better resemble the three-dimensional subcortical actin/spectrin meshwork found in erythrocytes, where structural integrity is balanced with mechanical deformability. The slightly extended longitudinal spacing of Palm3 (∼143 nm) compared to its circumferential distribution (∼132 nm) is consistent with the observations in neuronal axons, where microtubular stretch regulates β-Spectrin organization and supports its full-length extension (Zhong *et al*, 2014). In line with this hypothesis, the longitudinal microtubule network of auditory hair cells is primarily oriented in an apico-basal direction (Steyger *et al*, 1989; Furness *et al*, 1990; Voorn *et al*, 2024), and would thus enable a greater longitudinal spacing of Palm3. Additionally, Palm3 exhibited enrichment peaks along the OHC circumference that overlapped with those of ɑII- and βI- Spectrin. Together with the positive ICQ values, this finding suggests tight colocalization within the same subcellular compartment and implies a functional dependency.

### Palm3 stabilizes the submembrane cytoskeleton to maintain SSC integrity

As prominent structures of the OHC lateral wall (Dieler *et al*, 1991), SSCs have been proposed to support electromotility by modulating voltage signaling – either via electro-osmotic effects near the plasma membrane (Kachar *et al*, 1986) or by facilitating rapid transmission of membrane potential changes to Prestin-rich domains (Song & Santos-Sacchi, 2015). Moreover, SSCs facilitate vesicular trafficking (Kaneko *et al*, 2006) and likely serve as intracellular calcium reservoirs (Schulte, 1993). In *Palm3*-KO OHCs, SSCs were strikingly reduced in number and displayed a vastly simplified complexity, with the few remaining structures appearing flattened and displaced from the plasma membrane. On first sight, this suggests intrinsic structural collapse of SSCs; however, their distal positioning more likely reflects secondary consequences of cytoskeletal disruption, potentially due to impaired anchoring to the submembrane cytoskeleton in the absence of Palm3. Considering the reduced spectrin expression in *Palm3*-KO OHCs, this anchoring defect is more plausibly a result of a broader collapse of the submembrane cytoskeleton, rather than being a direct consequence of *Palm3* loss alone. Collectively, these results indicate that Palm3 plays a critical role in the maintenance of the SSCs, likely by contributing to the stability of the submembrane cytoskeleton. Interestingly, prior research on salicylate-induced SSC damage has linked SSC disruption to decreased stiffness of the lateral wall, leading to compromised OHC lateral wall mechanics and impaired electromotility (Lue & Brownell, 1999; Song & Santos-Sacchi, 2015; Shehata *et al*, 1991). Such functional deficits may underlie the near-complete loss of cochlear amplification observed at three weeks of age, even in the presence of surviving OHCs.

### Palm3 as a component of the pillar complex?

‘Pillars’ have been described as electron-dense, periodically-spaced structures that connect the OHC plasma membrane to the underlying submembrane cytoskeleton. Early ultrastructural studies reported pillars to average around ∼30 nm in length (Holley & Ashmore, 1990). Further investigations revealed that pillars likely behave as entropic springs, varying in length from 17 to 51 nm, with an average spacing of 26 ± 11 nm along actin filaments. Moreover, pillars were found to extend into a Y-shape as the distance between the OHC plasma membrane and the underlying cytoskeleton increases, and flatten against the membrane as that distance decreases (Forge, 1991; Holley *et al*, 1992; Triffo *et al*, 2019). Despite their discovery in the late 1980s, the molecular identity of these structures remains unresolved to date. We propose that Palm3 may serve as a key component of the pillar protein complexes. Several lines of evidence support this hypothesis: (i) Palm3 contains a CaaX motif, confirming its definitive localization to the plasma membrane. (ii) Pillar-like structures are present in both OHCs and IHCs (Furness & Hackney, 1990), which is consistent with Palm3 expression in both cell types. (iii) Due to the overlapping localization pattern of Palm3/spectrins, the dramatic reduction of spectrins within the lateral wall of *Palm3*-KO OHCs and drawing from the analogy with Palm1 (i.e., assuming that spectrin-binding is conserved across paralemmins), Palm3 may similarly associate with spectrins, although direct evidence to substantiate this link remains to be established. (iv) Palm3 exhibits a seemingly periodic and clustered distribution that resembles the spectrin staining pattern in erythrocytes, suggesting a potential structural parallel in cytoskeletal organization, which however differs in the extent of spectrin extension. (vi) Finally, the flexible, entropic spring-like behavior of pillars is compatible with the predicted intrinsically-disordered nature of Palm3, which could allow Palm3 to accommodate dynamic changes in cytoskeletal architecture along the OHC lateral wall.

However, certain discrepancies emerge when comparing the known dimensions and periodicity of pillars with our Palm3 data. Specifically, Palm3 puncta observed through 2D-STED NND and line profile-based peak-to-peak analyses show significantly wider spacing (∼125-194 nm) than the reported EM-based spacing range of ∼17-51 nm for individual pillars. This difference may stem from technical limitations of the immunofluorescence techniques, including reduced spatial accuracy due to bulky antibody-fluorophore conjugates and potential clustering artifacts induced by chemical fixation. Importantly, variations in sample preparation, particularly the use of distinct chemical fixation procedures must be considered, as they can introduce significant variability. Alternatively, it also remains a possibility that Palm3 is not part of the pillar complexes but instead contributes more broadly to lateral membrane stability, which is a possibility that warrants further exploration through high-resolution structural and biochemical approaches. In conclusion, our data suggests that Palm3 represents a formidable – and, to date, to our knowledge the *only* – protein candidate to contribute to pillar complexes since these structures were first described.

### Viral rescue at early postnatal stages can successfully correct *Palm3* loss from OHCs

Given the early onset of OHC degeneration in *Palm3*-KO mice and recent associations between *Palm3* mutations and age-related hearing loss in humans (Cornejo-Sanchez *et al*, 2025), our evaluation of AAV-mediated *Palm3* rescue reveals important insights into its potential for future therapeutic intervention. Since OHC degeneration in *Palm3-*KO mice begins around hearing onset, this period represents a critical early postnatal window of opportunity, during which gene therapy may effectively counteract the pathological consequence of *Palm3* loss. In line with this, our study shows that postnatal reintroduction of wildtype *Palm3* can alleviate the detrimental phenotype associated with its deficiency. Specifically, OHC survival was markedly improved compared to the pronounced degeneration seen in *Palm3*-KO mice by five weeks of age. In addition to enhanced survival, *Palm3*-rescued OHCs exhibited notable restoration in cell length, indicating a clear recovery toward normal cellular morphology. At the molecular level, *Palm3* reintroduction restored proper expression of ɑII-Spectrin and corrected the aberrant distribution of Prestin along the OHC lateral membrane. These findings demonstrate that postnatal re-introduction of *Palm3* effectively reverses the key deficits associated with its loss and further emphasize the essential role of Palm3 in stabilizing the lateral membrane and maintaining OHC structural integrity.

However, despite the significant cellular and morphological improvements observed in *Palm3*-rescued OHCs, these structural restorations translated into only modest recovery of hearing function, pointing to several limitations that may have collectively constrained the therapeutic outcome: (i) The transduction efficiency was highly variable across rescue animals, with especially poor targeting of vulnerable OHCs and more consistent delivery to the less affected IHCs. (ii) Off-target effects due to the broad tropism of the AAV.S vector (Ivanchenko *et al*, 2021) – including *Palm3* overexpression in IHCs, stereociliar mislocalization that may exert a negative impact on mechanotransduction, and ectopic transduction of SGNs – likely compromised cellular structural homeostasis, introduced cellular stress, and impaired synaptic function to ultimately further compromise auditory function. Lastly, (iii) potential AAV-induced cytotoxicity (Meyers *et al*, 2001; Hirsch *et al*, 2011; Rapti *et al*, 2015; Johnston *et al*, 2021), which has previously also been reported in the auditory system (Wei *et al*, 2025), cannot be excluded as a contributing factor to IHC loss. Future studies should aim for improved vector specificity, tightly controlled gene expression, and optimized delivery strategies to enhance *Palm3* gene therapy in future studies. Nevertheless, despite current limitations, this study provides a proof-of-concept which lays the groundwork for advancing the therapeutic potential of *Palm3*-based gene therapy.

## Concluding remarks

In summary, this study identifies Palm3 as a critical molecular component of the peripheral auditory system, that is essential for maintaining the structural integrity of the OHC lateral wall. Our findings introduce a promising novel deafness gene candidate and address a major gap in the molecular understanding of cochlear architecture – the molecular nature of the elusive pillar complexes between the plasma membrane and the cortical cytoskeleton.

## Acknowledgements

We would like to express our utmost gratitude to Christiane Senger-Freitag, Sandra Gerke and Jana Erlmoser for expert technical support, Nicole Metzendorf for initial AAV plasmid cloning and Rudolph Glückert for assistance with the ultrathin melamine-sectioning. This work was funded by a research grant from the Deutsche Forschungsgemeinschaft (DFG, German Research Foundation) – Project Number VO 2466/1-1 to CV and MWK, an *Otto Creutzfeldt Fellowship* of the Elisabeth and Helmuth Uhl Foundation (to CV), and intramural funding of the Medical University Innsbruck (to CV) and the CRC 889/3 project A07 to CW. The High-pressure-freezer was funded by the Deutsche Forschungsgemeinschaft (DFG, German Research Foundation) – Project number 257917528. The transmission electron microscope JEM1011 was funded by the Deutsche Forschungsgemeinschaft (DFG, German Research Foundation) – Project number 130725592. The transmission electron microscope JEM2100Plus was funded by REACT-EU from “Europäische Fonds für regionale Entwicklung (EFRE)” with the project *“ETomoH&H: Elektronentomografie an Herz und Hirn”* and the University Medical Center Göttingen.

## Author contributions

CV and MWK conceived the project and designed the experiments. VCH and CV performed immunohistochemistry, confocal imaging and data analysis. VCH performed STED immunohistochemistry, melamine sample embedding and STED imaging. VCH did chemical clearing and lightsheet microscopy. DD, MFK and DO performed OHC electrophysiology and analysis. GH performed cDNA sequencing, cloning and purification of Palm3 fusion proteins for immunisation, affinity-purification and pre-testing (Western blot) of sera. IB and NS measured and analyzed ABRs and DPOAEs. LS performed HPF/FS and TEM with assistance from VCH. CU and CW performed the electron tomography. KK designed and generated rescue AAVs. VCH, CV and MWK wrote the paper, VCH and CV generated all Figures. All co-authors revised the manuscript.

