## Supplementary materials for "Paralemmin-3 sustains the integrity of the lateral plasma membrane and subsurface cisternae of auditory hair cells"

#### Supplemental Figures

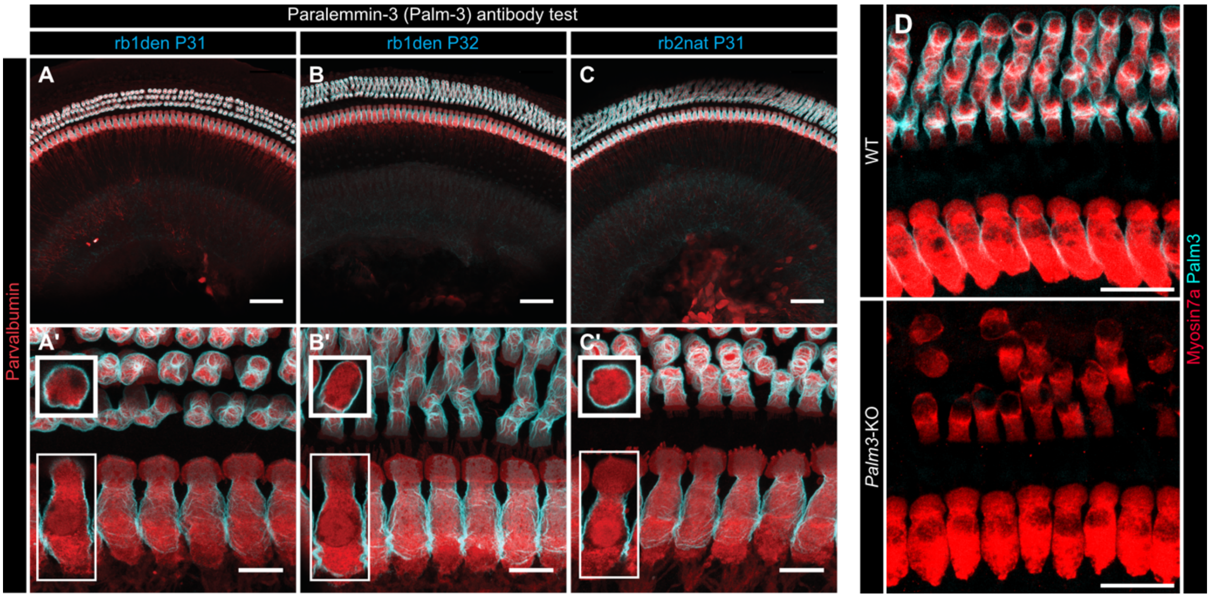


**Supplementary figure 1. Palm3 IF staining specificity controls: Three independent custom-made Palm3 antibodies produced the same patterns in murine IHC and OHC lateral plasma membranes (A-C), which were abolished in Palm3-KO mouse tissue (D).**

(A-C) Representative confocal maximum projection of z-stacks showing organs of Corti overviews from P15 WT mice stained with three different Palm3 antibodies: (A) anti-Palm3 p31 rb1 den, (B) anti-Palm3 p32 rb1 den (N-terminal & C-terminal denatured immunogens), and (C) anti-Palm3 p31 rb2 nat (N-terminal native immunogen). Hair cells are immunostained against Palm3 (cyan) and Myosin7A (red). Scale bars: 50 µm.

(A’-C’) High magnification maximum projections of organs of Corti for all three Palm3 antibodies from A-C. Insets show representative single optical sections of individual IHC and OHC for each AB to illustrate the exclusive lateral plasma membrane localization of Palm3. Hair cells are immunostained for Palm3 (cyan) and Myosin7A (red). Scale bars: 10 µm.

(D) Representative confocal maximum z-projections of apical turn whole-mount organs of Corti from WT and *Palm3*-KO at P35. OHCs were labeled against the hair cell marker Myosin7a (red) and Palm3 (cyan). Scale bars: 20 µm.

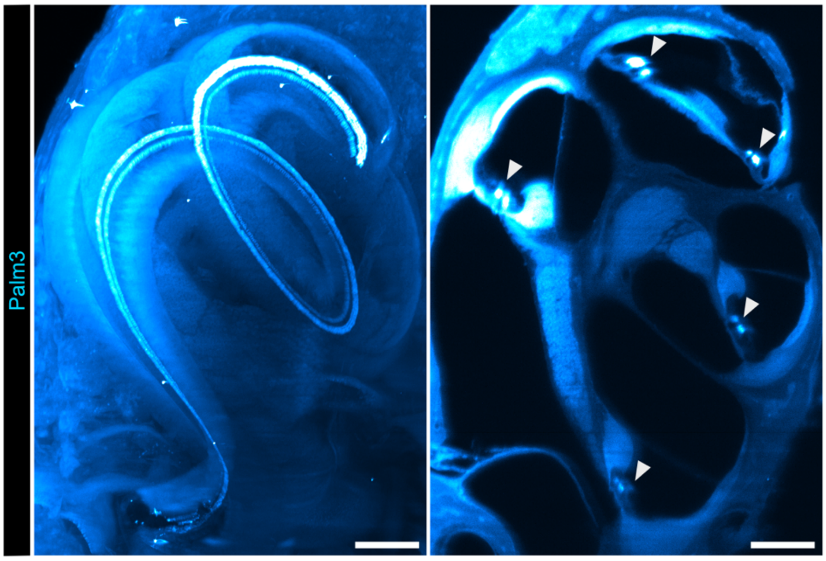


**Supplementary figure 2. Unprocessed lightsheet microscopy image of Palm3-stained, iDisco-cleared WT cochlea.**

Representative maximum z-projection (left panel) and single optical section (right panel) of lightsheet microscopy on iDisco-cleared cochlea of a P14 WT mouse. Hair cells are stained with Palm3 (cyan hot). Filled arrowheads on the right panel indicate rows of hair cells within the chemically-cleared cochlea. Scale bars: 100 µm. The lightsheet image from **Figure 1** was processed in *Imaris* 10.2.0 to exclude auto-fluorescent signals from surrounding cochlear bone and enhance clarity of Palm3 staining. First, 3D reconstructions of Palm3 immunofluorescence were rendered using the *surfaces* algorithm with the surface size set to 0.65 µm. Thresholding was applied to exclude artifacts based on quality of the staining. A clipping mask specific to Palm3 immunofluorescence was generated by manually removing surfaces rendered outside of Palm3 immunofluorescence signal. The masked immunofluorescent signal was then exported to ImageJ/Fiji software (Schneider *et al*, 2012) to generate a maximum projection of z-stack.

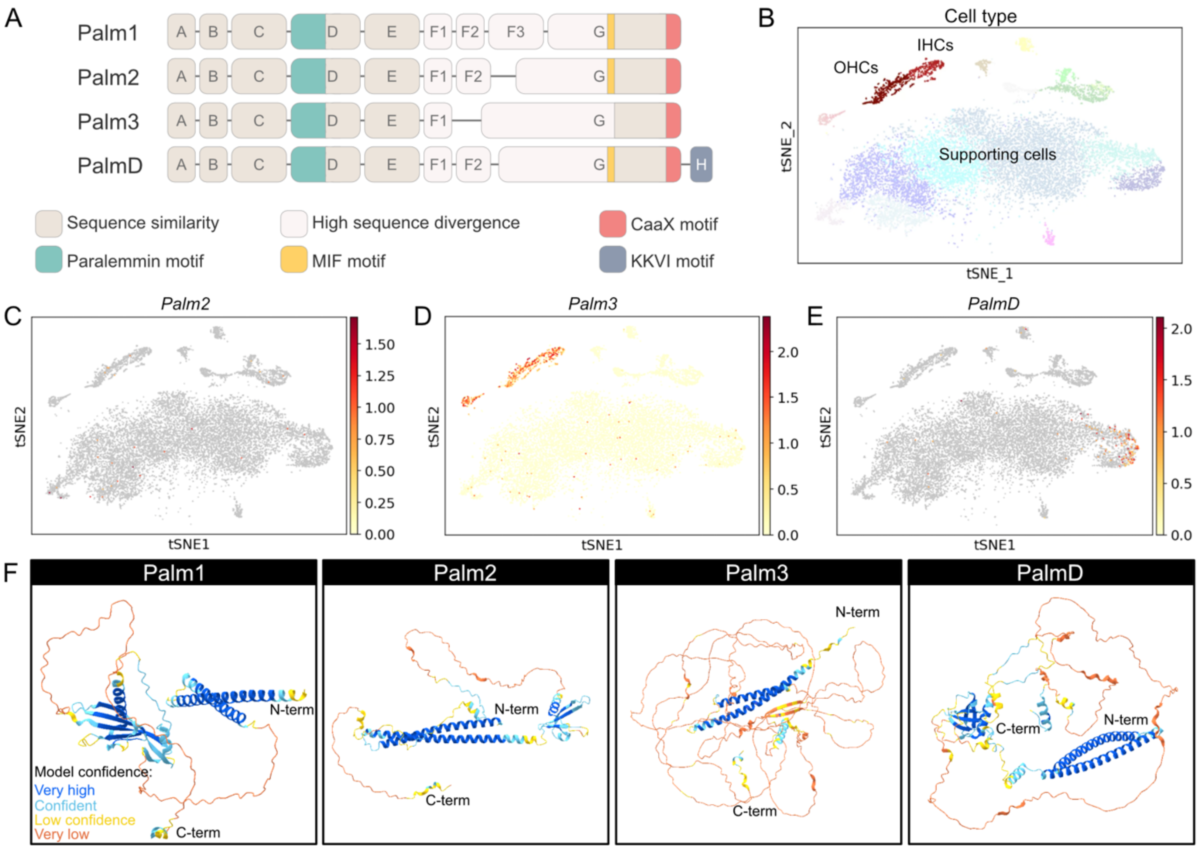


**Supplementary figure 3. The Paralemmin protein family.**

(A) Schematic representation of conserved organization among mammalian paralemmin isoforms. Human Palm1 is composed of 389 amino acids (AA), Palm2 of 381 AA, Palm3 of 675 AA, and Palmd of 552 AA. Boxes represent the paralogous exons of paralemmin isoforms. Modified from Hultqvist et al., 2012.

(B) A t-SNE plot showing cell clustering in the organ of Corti based on feature similarities. Each point represents a single cell, with clusters formed based on shared features. Colors indicate different cell types found in the murine organ of Corti, provided with a simplified visualization of distinct cell populations to aid more accessible interpretations.

(C-E) t-SNE plots displaying scRNAseq data on the expression of (C) *Palm2*, (D) *Palm3*, (E) *Palmd* in the murine organ of Corti. No scRNA-seq data for Palm1 is available on gEAR. Colors represent expression fold change or transcript abundance, with warmer colors indicating transcript abundance and cooler colors reflecting lower amounts, as shown in the accompanying heatmap. (Orvis *et al*, 2021; https://umgear.org/).

(F) Three-dimensional (3D) structural prediction of paralemmin protein isoforms found in *Mus musculus* (mouse). In mice, Palm1 is composed of 385 amino acids (AA), Palm2 of 378 AA, Palm3 of 736 AA, and Palmd of 553 AA. Model confidence scores are color-coded along the amino acid sequences, with higher confidence regions highlighted in cooler colors and lower confidence regions in warmer colors. Structures were generated using Alphafold (https://alphafold.ebi.ac.uk).

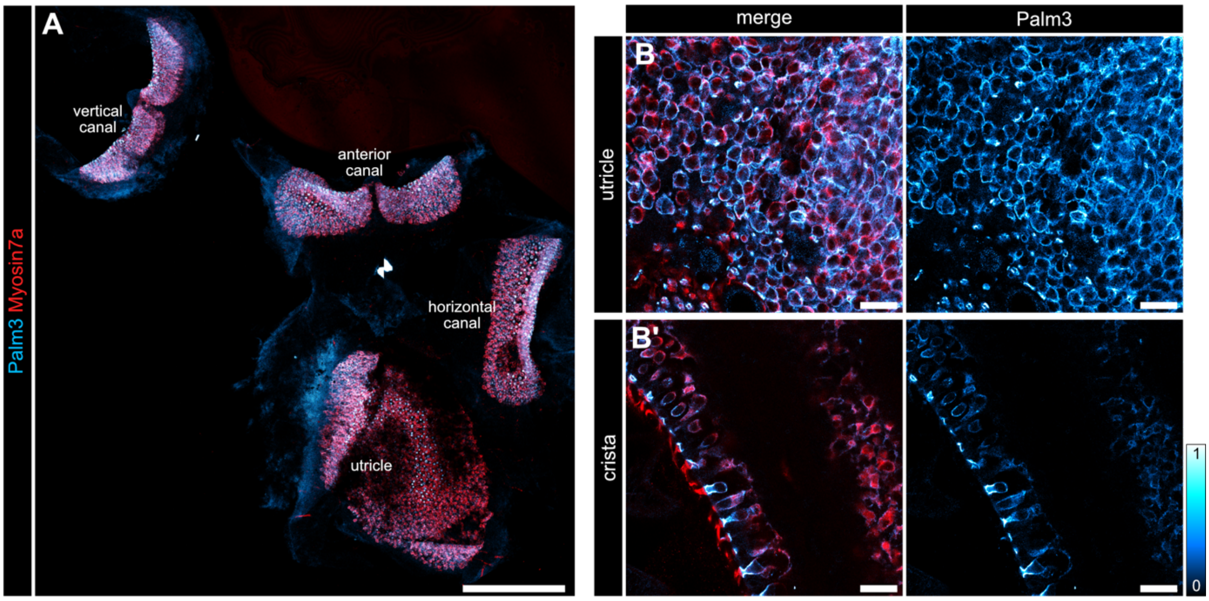


**Supplementary figure 4. Palm3 is expressed in hair cells of the murine vestibular system.**

(A) Representative confocal maximum projection of z-stacks showing an overview of the vestibular system of a P48 WT mouse stained against Palm3 (cyan hot) and Myosin7a as a context marker (red). Scale bar: 250 µm.

(B,B') Single optical section of (B) the utricle and (B') a crista of the vestibular system at higher magnification. Scale bars: 20 µm.

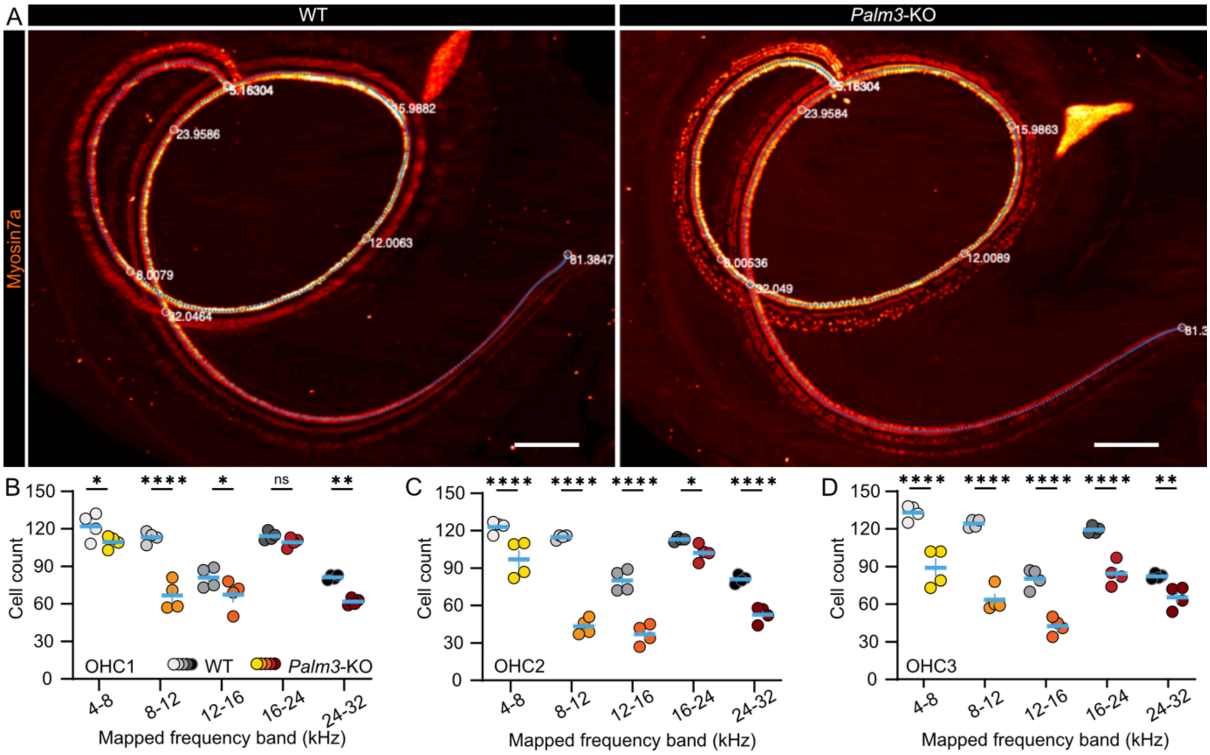


**Supplementary figure 5. Genetic loss of Palm3 leads to OHC degeneration along the entire tonotopic axis.**

(A) Representative maximum z-projections of lightsheet microscopic images acquired from iDisco-cleared WT and *Palm3*-KO cochleae around hearing onset (P14). Hair cells were stained against Myosin7a as a hair cell specific marker. Tonotopic mapping of hair cells was based on Greenwood's function (Greenwood, 1961) with previously described parameters (Müller *et al*, 2005). Blue dotted lines indicate 500 reference points that were interpolated along the cochlear spiral based on the frequency-mapping algorithm. Approximate frequency regions are labeled and given in kHz. Scale bars: 100 µm. (B-D) Single value plots of WT (grey) and *Palm3*-KO OHC count show varying degrees of degeneration in the indicated frequency regions for (B) OHC row 1 (innermost), (C) OHC row 2 (middle), and (D) OHC row 3 (outermost). $N_{animals}$= 2, $n_{Cochlea}$ = 4 for WT and $N_{animals}$= 2, $n_{Cochlea}$ = 4 for *Palm3*-KOs.

Data information: Two-way ANOVA with Holm-Šídák‘s multiple comparisons correction: ****$P$ <0.0001; Mean ± SEM.

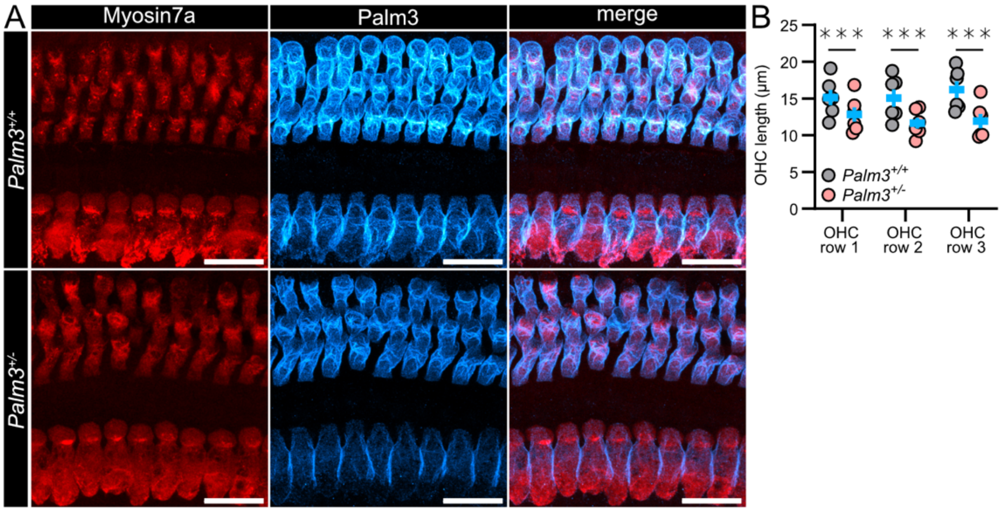


**Supplementary figure 6. Attenuated expression and OHC length in Palm3$^{+/-}$.**

(A) Representative confocal maximum z- projections of OHCs stained against Myosin7a (red) and Palm3 (cyan hot) from 8-week-old WT and *Palm3*$^{+/-}$.

(B) *Palm3*$^{+/-}$ OHCs exhibit reduced cell length in all 3 OHC rows. Two-way ANOVA with Holm-Šídák's multiple comparisons correction: **** P < 0.0001; Mean ± SEM.

Data information: $N_{animals}$= 3, $n_{Corti}$ = 6 each for WT and *Palm3*$^{+/-}$

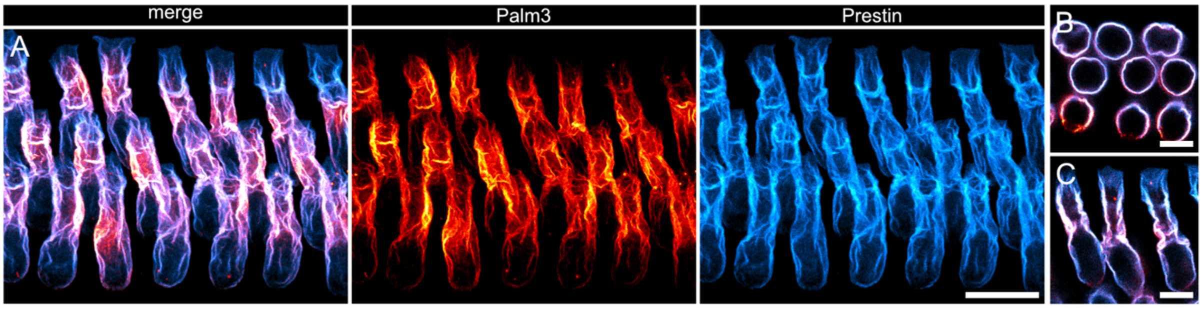


**Supplementary figure 7. Palm3 colocalizes with Prestin in the OHC lateral plasma membrane.** (A) Representative confocal maximum projection of z-stacks showing organs of Corti from P14 WT mouse. OHCs are stained against Palm3 (red) and Prestin (cyan). Scale bars: 10 µm. (B-C) Single optical sections showing (B) circumferential and (C) longitudinal distribution of Palm3 (red) and Prestin (cyan). Scale bars: 5 µm.

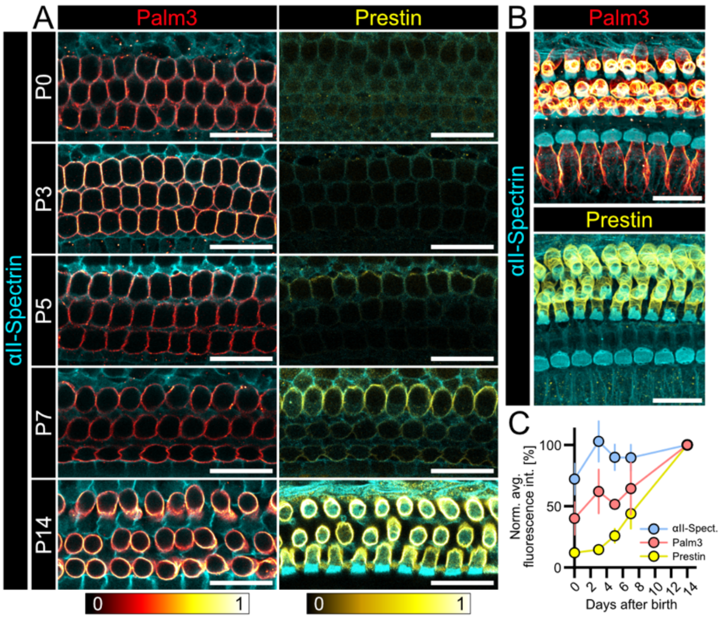


**Supplementary figure 8. Palm3 precedes Prestin arrival in the OHC lateral membrane.**

(A) OHCs were stained against either Palm3 (red hot, left panels) or Prestin (yellow hot, right panels), each in combination with αII-Spectrin (cyan) as context. Representative confocal single optical sections of apical organs of Corti from P0, P3, P5, P7 and P14 *Palm3*-WT animals. All samples were taken from comparable tonotopic regions, imaged and subsequently processed in parallel. Scale bars: 20 µm.

(B) Representative maximum z-projection of whole-mount apical organs of Corti from *Palm3*-WT animals at P14. Scale bars: 20 µm.

(C) Normalized average fluorescence intensity of αII-Spectrin-, Palm3-, and Prestin-staining plotted against age (days after birth) shows αII-Spectrin and Palm3 to precede arrival of Prestin in the OHC lateral membrane. For Palm3 and Prestin: $N_{animals}$= 4, $n_{Corti}$ = 4 for P0; $N_{animals}$= 4, $n_{Corti}$ = 4 for P3; $N_{animals}$= 4, $n_{Corti}$ = 4 for P5; $N_{animals}$= 4; $n_{Corti}$ = 4 for P7; and $N_{animals}$= 4, $n_{Corti}$ = 4 for P14. For αII-Spectrin: $N_{animals}$= 4, $n_{Corti}$ = 8 for P0; $N_{animals}$= 4, $n_{Corti}$ = 8 for P3; $N_{animals}$= 4, $n_{Corti}$ = 8 for P5; $N_{animals}$= 4, $n_{Corti}$ = 8 for P7; and $N_{animals}$= 4, $n_{Corti}$ = 8 for P14. Mean ± SEM.

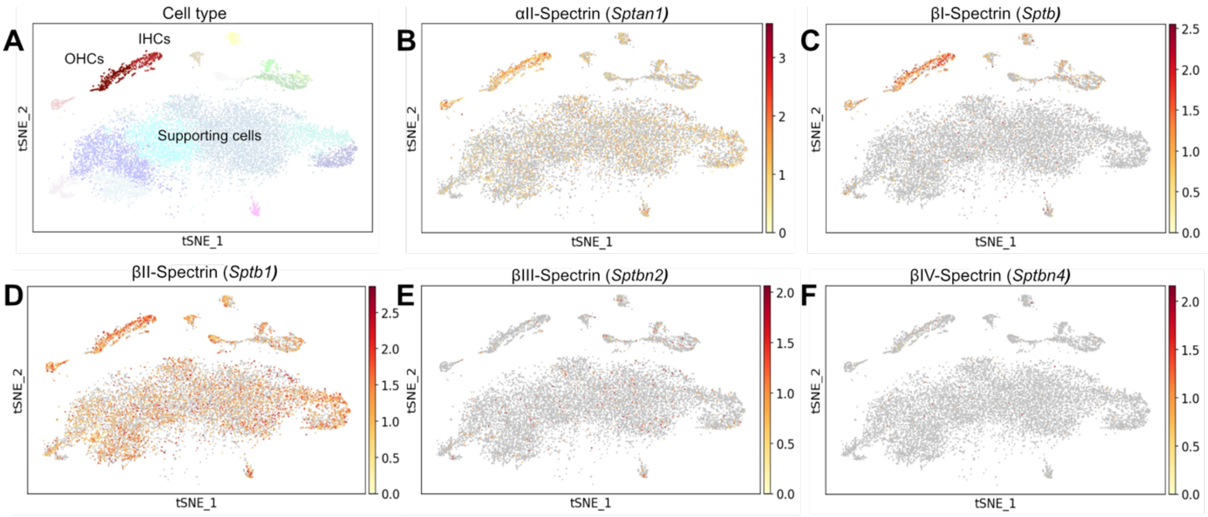


**Supplementary figure 9. Single cell RNA sequencing (scRNA-seq) t-distributed Stochastic Neighbor Embedding (t-SNE) plots of spectrin isoform expression in the organ of Corti of P1 mice.**

(A) A t-SNE plot showing cell clustering in the organ of Corti based on feature similarities. Each point represents a single cell, with clusters formed based on shared features. Colors indicate different cell types found in the organ of Corti, provided with a simplified visualization of distinct cell populations to aid more accessible interpretations.

(B-F) t-SNE plots displaying scRNAseq data on the expression of (B) αII-Spectrin, (C) βI-Spectrin, (D) βII-Spectrin, (E) βIII-Spectrin, and (F) βIV-Spectrin in the organ of Corti. No scRNA-seq data for erythrocytic αI-Spectrin and non-erythrocytic βV-Spectrin are available on gEAR. Colors represent expression fold change or transcript abundance, with warmer colors indicating transcript abundance and cooler colors reflecting lower amounts, as shown in the accompanying heatmap. (Orvis *et al*, 2021; https://umgear.org/)

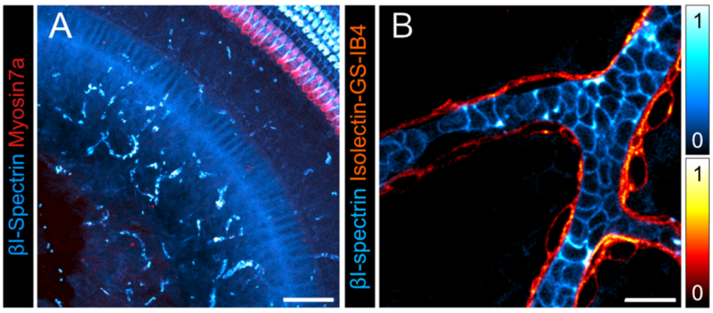


**Supplementary figure 10. βI-Spectrin IF staining is observed in erythrocytes within the organ of Corti.**

(A) Representative confocal maximum projection of z-stacks showing organs of Corti from a P19 WT mouse stained with βI-Spectrin (cyan hot) and Myosin7a as a context marker (red). Scale bar: 50 µm. (B) Magnified view of a single blood vessel filled with βI-Spectrin-positive erythrocytes (cyan hot), counterstained with Isolectin-GS-IB4 to visualize the endothelial lining (red hot). Scale bar: 10 µm.

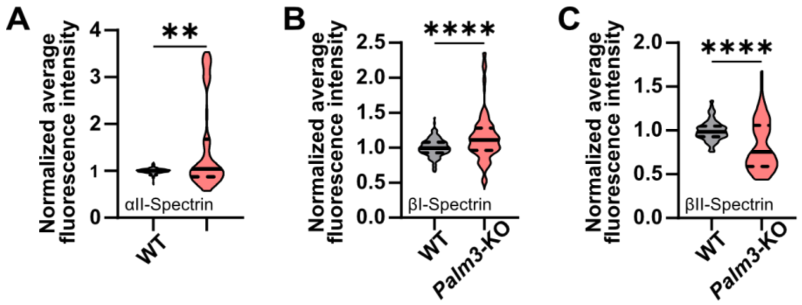


**Supplementary figure 11. Effect of Palm3 absence on the expression of spectrin isoforms in OHC cuticular plates.**

(A-C) Violin plots showing average fluorescence intensity of (A) αII-Spectrin ($N_{animals}$= 5, $n_{Corti}$ = 10, $n_{OHC}$ = 265 for for WT and $N_{animals}$= 5, $n_{Corti}$ = 10, $n_{OHC}$ = 175 for *Palm3*-KO), (B) βI-Spectrin ($N_{animals}$= 5, $n_{Corti}$ = 10, $n_{OHC}$ = 156 for WT and $N_{animals}$= 5, $n_{Corti}$ = 10, $n_{OHC}$ = 125 for *Palm3*-KO) and (C) βII-Spectrin ($N_{animals}$= 3, $n_{Corti}$ = 6, $n_{OHC}$ = 112 for WT and $N_{animals}$= 3, $n_{Corti}$ = 6, $n_{OHC}$ = 77 for *Palm3*-KO) in the cuticular plates of WT and *Palm3*-KO OHCs. Two-tailed Mann-Whitney test: ** P < 0.01; **** P < 0.0001.

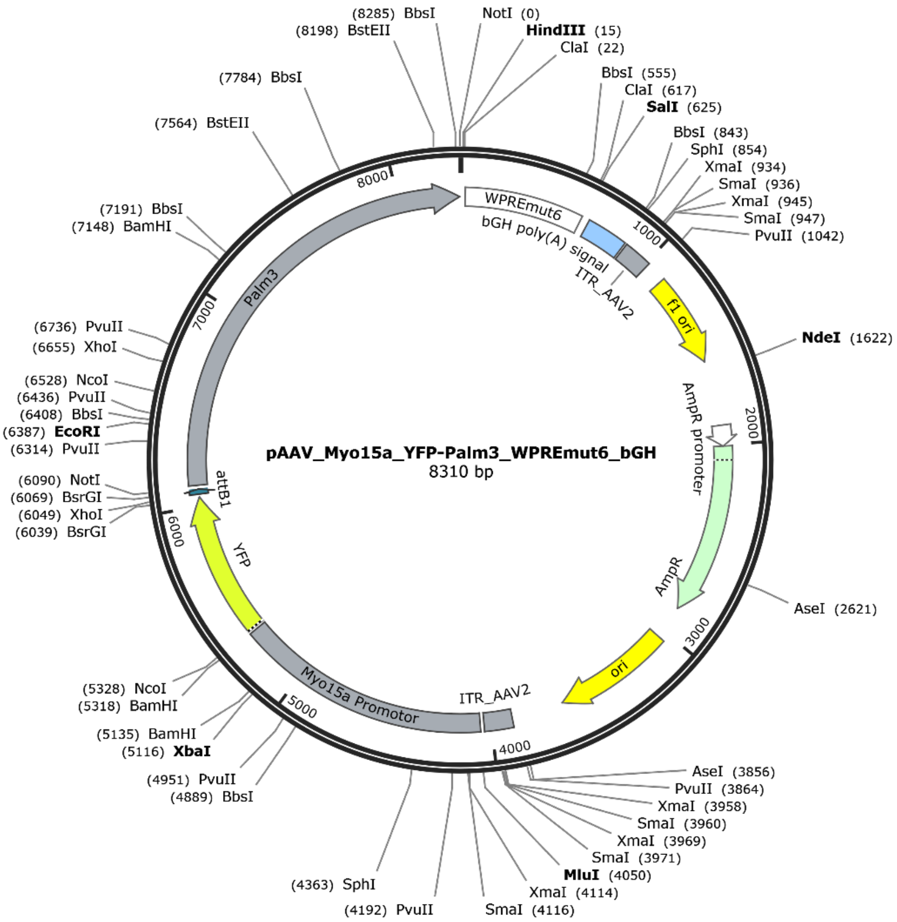


**Supplementary figure 12. AAV.S_Myo15a_YFP-Palm3**

The vector, constructed on a pAAV backbone, encodes mouse Palm3 fused at its N-terminus with yellow fluorescent protein (YFP). The fusion cDNA is inserted between the human *Myo15a* promoter and a non-oncogenic variant of the Woodchuck hepatitis virus posttranscriptional regulatory element (*WPRE*), followed by bovine growth hormone (*bGH*) polyadenylation signal sequences. The entire cassette flanked by inverted terminal repeats (*ITRs*).

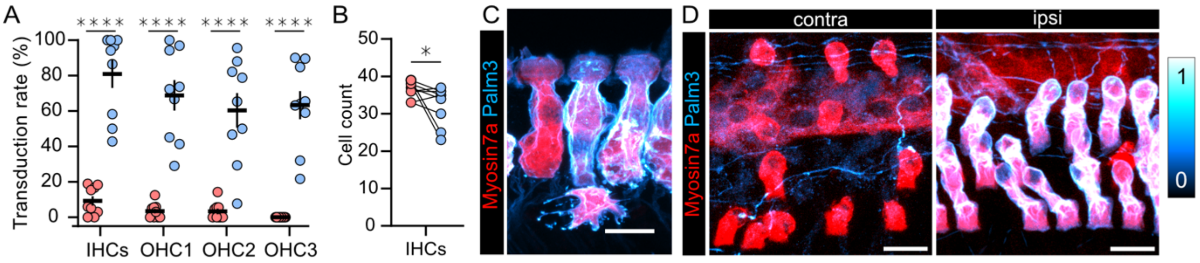


**Supplementary figure 13. Limitations of AAV-mediated *Palm3*-rescue**

(A) High variability in the transduction efficiency across individual animals, shown by scattered single value plots quantifying viral transduction rates. Two-way ANOVA with Holm-Šídák's multiple comparisons correction: **** $P < 0.0001$.

(B) Single value plots showing IHCs counts within a 300 µm x 300 µm region of interest to be slightly declined in *Palm3*-rescue ears. Connecting lines identify individual animals. Two-tailed Wilcoxon matched-pairs signed rank test: * $P < 0.05$.

(C) *Palm3*-transduced IHC displaying aberrant cell shape. Hair cells stained for Myosin7a (red) and Palm3-YFP (cyan hot). Scale bar: 10 µm.

(D) Representative confocal maximum z-projections of OHCs of contralateral- and ipsilateral ears of 5-week-old *Palm3*-KO animals. Note the ectopic expression of Palm3-YFP in SGN fibers. Cortis stained for Myosin7a (red) and Palm3-YFP (cyan hot). Scale bars: 10 µm.

### Supplemental Tables

#### Supplementary Table 1: Statistics summary I

| | | | WT | | *Palm3* -KO | | Statistical test | F value | *P* value | Significance |
|---|---|---|---|---|---|---|---|---|---|---|
| | | | Mean ± SEM | SD | Mean ± SEM | SD | | | | |
| ABR Threshold | 3 weeks | 4 kHz | 57.8 ± 2.78 | 8.33 | 86.1 ± 1.62 | 4.86 | Two-way ANOVA with Holm-Šídák's multiple comparisons correction | F (1,135) = 291.0 | <0.0001 | **** |
| | | 6 kHz | 41.1 ± 3.61 | 10.8 | 67.8 ± 2.22 | 6.67 | | | <0.0001 | **** |
| | | 8 kHz | 34.4 ± 2.94 | 8.82 | 59.4 ± 2.94 | 8.82 | | | <0.0001 | **** |
| | | 12 kHz | 35.6 ± 3.77 | 11.3 | 51.1 ± 2.61 | 7.82 | | | <0.0001 | **** |
| | | 16 kHz | 39.4 ± 2.69 | 8.08 | 52.2 ± 2.22 | 6.67 | | | <0.0001 | **** |
| | | 24 kHz | 42.8 ± 2.78 | 8.33 | 72.8 ± 3.45 | 10.3 | | | <0.0001 | **** |
| | | 32 kHz | 51.7 ± 4.08 | 12.3 | 82.8 ± 2.78 | 8.33 | | | <0.0001 | **** |
| | | click | 30.0 ± 0.00 | 0.00 | 65.6 ± 1.76 | 5.27 | | | <0.0001 | **** |
| | 11-12 weeks | 4 kHz | 70.6 ± 2.39 | 6.78 | 90.0 ± 0.00 | 0.00 | | F (1,117) = 828.5 | <0.0001 | **** |
| | | 6 kHz | 49.4 ± 1.99 | 5.63 | 86.9 ± 1.32 | 3.72 | | | <0.0001 | **** |
| | | 8 kHz | 45.0 ± 1.89 | 5.35 | 85.0 ± 1.34 | 3.78 | | | <0.0001 | **** |
| | | 12 kHz | 41.9 ± 1.88 | 5.30 | 86.3 ± 1.57 | 4.43 | | | <0.0001 | **** |
| | | 16 kHz | 46.9 ± 1.62 | 4.58 | 89.4 ± 0.63 | 1.77 | | | <0.0001 | **** |
| | | 24 kHz | 42.9 ± 1.49 | 3.93 | 90.0 ± 0.00 | 0.00 | | | <0.0001 | **** |
| | | 32 kHz | 60.0 ± 3.62 | 9.57 | 90.0 ± 0.00 | 0.00 | | | <0.0001 | **** |
| | | click | 31.3 ± 1.25 | 3.54 | 97.5 ± 1.64 | 4.63 | | | <0.0001 | **** |
| DPOAE Intensity | 3 weeks | 6 kHz | -14.3 ± 1.87 | 5.29 | -18.9 ± 1.99 | 5.96 | Two-way ANOVA with Holm-Šídák's multiple comparisons correction | F (3,242) = 68.2 | <0.0001 | **** |
| | | 8 kHz | -9.51 ± 1.48 | 4.19 | -14.0 ± 3.12 | 9.36 | | | <0.0001 | **** |
| | | 11 kHz | -0.04 ± 2.15 | 6.08 | -17.9 ± 2.09 | 6.29 | | | <0.0001 | **** |
| | | 16 kHz | 3.31 ± 3.09 | 8.76 | -16.9 ± 2.87 | 8.62 | | | <0.0001 | **** |
| | | 23 kHz | 0.91 ± 3.73 | 10.6 | -21.2 ± 2.01 | 6.02 | | | <0.0001 | **** |
| | | 32 kHz | -8.51 ± 3.61 | 10.2 | -20.6 ± 2.18 | 6.53 | | | <0.0001 | **** |
| | | 45 kHz | -19.1 ± 1.74 | 4.91 | -23.8 ± 0.59 | 1.79 | | | <0.0001 | **** |
| | 11-12 weeks | 6 kHz | -19.7 ± 1.67 | 4.42 | -17.2 ± 3.42 | 9.67 | | F (3,200) = 20.9 | <0.0001 | **** |
| | | 8 kHz | -10.9 ± 2.87 | 7.60 | -19.9 ± 1.79 | 5.07 | | | <0.0001 | **** |
| | | 11 kHz | -6.85 ± 5.13 | 13.6 | -22.3 ± 1.36 | 3.83 | | | <0.0001 | **** |
| | | 16 kHz | 4.03 ± 2.43 | 6.42 | -18.3 ± 2.60 | 7.35 | | | <0.0001 | **** |
| | | 23 kHz | -5.05 ± 3.71 | 9.82 | -22.7 ± 0.82 | 2.31 | | | <0.0001 | **** |
| | | 32 kHz | -19.7 ± 2.89 | 7.66 | -23.3 ± 0.42 | 1.17 | | | <0.0001 | **** |
| | | 45 kHz | -23.8 ± 0.85 | 2.24 | -21.3 ± 3.11 | 8.79 | | | <0.0001 | **** |
| Age-dependent Degeneration | IHC | 2-3 weeks | 24.8 ± 0.86 | 2.44 | 23.4 ± 0.65 | 1.85 | Two-way ANOVA with Holm-Šídák's multiple comparisons correction | F (1,54) = 13.5 | 0.0022 | ** |
| | | 7-8 weeks | 25.0 ± 1.16 | 2.83 | 23.5 ± 0.22 | 0.55 | | | 0.0022 | ** |
| | | 25 weeks | 24.5 ± 0.63 | 1.77 | 23.1 ± 0.46 | 1.22 | | | 0.0022 | ** |
| | | 50 weeks | 23.0 ± 0.38 | 1.0 | 20.3 ± 0.77 | 2.19 | | | 0.0022 | ** |
| | OHC1 | 2-3 weeks | 25.8 ± 0.56 | 1.58 | 21.9 ± 1.33 | 3.76 | | F (1,54) = 118.0 | <0.0001 | **** |
| | | 7-8 weeks | 25.7 ± 0.56 | 1.37 | 17.5 ± 1.23 | 3.02 | | | <0.0001 | **** |
| | | 25 weeks | 25.9 ± 0.48 | 1.36 | 9.71 ± 1.48 | 3.90 | | | <0.0001 | **** |
| | | 50 weeks | 23.3 ± 0.37 | 1.04 | 5.50 ± 1.50 | 4.24 | | | <0.0001 | **** |
| | OHC2 | 2-3 weeks | 25.9 ± 0.64 | 1.81 | 19.1 ± 1.25 | 3.52 | | F (1,54) = 146.1 | <0.0001 | **** |
| | | 7-8 weeks | 25.7 ± 1.02 | 2.50 | 18.5 ± 1.73 | 4.23 | | | <0.0001 | **** |
| | | 25 weeks | 25.8 ± 0.41 | 1.17 | 9.43 ± 0.97 | 2.57 | | | <0.0001 | **** |
| | | 50 weeks | 20.9 ± 1.33 | 3.76 | 2.63 ± 0.96 | 2.72 | | | <0.0001 | **** |
| | OHC3 | 2-3 weeks | 27.0 ± 0.53 | 1.51 | 18.6 ± 1.07 | 3.02 | | F (1,54) = 189.4 | <0.0001 | **** |
| | | 7-8 weeks | 25.2 ± 0.70 | 1.72 | 17.8 ± 1.40 | 3.43 | | | <0.0001 | **** |
| | | 25 weeks | 26.8 ± 0.56 | 1.58 | 8.71 ± 1.09 | 2.87 | | | <0.0001 | **** |
| | | 50 weeks | 21.1 ± 1.06 | 2.99 | 4.75 ± 1.21 | 3.41 | | | <0.0001 | **** |
| Lightsheet Cytocochleogram | OHC row 1 | 4-8 kHz | 122.0 ± 5.29 | 10.6 | 109.5 ± 2.39 | 4.79 | Two-way ANOVA with Holm-Šídák's multiple comparisons correction | F (4, 30) = 9.455 | 0.045 | * |
| | | 8-12 kHz | 113.3 ± 2.32 | 4.65 | 66.8 ± 5.72 | 11.44 | | | <0.0001 | **** |
| | | 12-16 kHz | 81.0 ± 4.08 | 8.17 | 67.3 ± 6.05 | 12.1 | | | 0.038 | * |
| | | 16-24 kHz | 114.3 ± 1.79 | 3.59 | 109.3 ± 1.93 | 3.86 | | | 0.34 | ns |
| | | 24-32 kHz | 81.3 ± 1.03 | 2.06 | 61.8 ± 1.38 | 2.75 | | | 0.003 | ** |
| | OHC row 2 | 4-8 kHz | 123.0 ± 2.42 | 4.83 | 97.0 ± 7.29 | 14.6 | | F (4, 30) = 20.19 | <0.0001 | **** |
| | | 8-12 kHz | 114.8 ± 0.95 | 1.89 | 43.3 ± 3.22 | 6.45 | | | <0.0001 | **** |
| | | 12-16 kHz | 80.0 ± 4.38 | 8.76 | 37.0 ± 4.06 | 8.12 | | | <0.0001 | **** |
| | | 16-24 kHz | 113.0 ± 0.82 | 1.63 | 102.3 ± 3.28 | 6.55 | | | 0.044 | * |
| | | 24-32 kHz | 81.0 ± 1.68 | 3.37 | 52.8 ± 3.19 | 6.39 | | | <0.0001 | **** |
| | OHC row 3 | 4-8 kHz | 133.0 ± 2.97 | 5.94 | 89.0 ± 7.61 | 15.2 | | F (4, 30) = 7.639 | <0.0001 | **** |
| | | 8-12 kHz | 124.3 ± 1.60 | 3.20 | 63.5 ± 4.87 | 9.75 | | | <0.0001 | **** |
| | | 12-16 kHz | 80.5 ± 3.93 | 7.85 | 42.5 ± 3.38 | 6.76 | | | <0.0001 | **** |
| | | 16-24 kHz | 119.3 ± 1.44 | 2.87 | 84.5 ± 4.77 | 9.54 | | | <0.0001 | **** |
| | | 24-32 kHz | 82.3 ± 1.11 | 2.22 | 65.5 ± 4.44 | 8.89 | | | 0.007 | ** |
| OHC Length | P11 | OHC row 1 | 15.8 ± 0.58 | 1.65 | 12.1 ± 0.63 | 1.79 | Two-way ANOVA with Holm-Šídák's multiple comparisons correction | F (1, 44) = 62.47 | <0.0001 | **** |
| | | OHC row 2 | 14.2 ± 0.53 | 1.49 | 11.2 ± 0.39 | 1.10 | | | <0.0001 | **** |
| | | OHC row 3 | 14.5 ± 0.49 | 1.40 | 11.4 ± 0.40 | 1.15 | | | <0.0001 | **** |
| | P21 | OHC row 1 | 14.9 ± 0.44 | 1.08 | 10.8 ± 0.32 | 0.78 | | F (1, 32) = 147.3 | <0.0001 | **** |
| | | OHC row 2 | 15.4 ± 0.23 | 0.57 | 9.48 ± 0.59 | 1.47 | | | <0.0001 | **** |
| | | OHC row 3 | 16.1 ± 0.64 | 1.58 | 7.52 ± 0.66 | 1.62 | | | <0.0001 | **** |
| NLC | Clin | P12-13 | 16.9 ± 0.41 | 0.99 | 7.33 ± 0.76 | 2.16 | Two-way ANOVA with Holm-Šídák's multiple comparisons correction | F (3,22) = 72.65 | <0.0001 | **** |
| | | P18-20 | 15.4 ± 0.56 | 1.84 | 6.92 ± 0.46 | 1.51 | | | <0.0001 | **** |
| | NLC | P12-13 | 6.25 ± 0.20 | 0.50 | 4.59 ± 0.17 | 0.49 | | F (3,22) = 26.6 | 0.175 | ns |
| | | P18-20 | 10.9 ± 0.98 | 3.26 | 4.34 ± 0.43 | 1.42 | | | <0.0001 | **** |
| | NLC/Clin | P12-13 | 0.32 ± 0.01 | 0.03 | 0.64 ± 0.05 | 0.14 | | F (3,22) = 6.63 | 0.0005 | *** |
| | | P18-20 | 0.53 ± 0.03 | 0.09 | 0.58 ± 0.05 | 0.17 | | | 0.465 | ns |
| | Vh | P12-13 | -26.6 ± 7.07 | 17.3 | -22.4 ± 2.06 | 5.81 | | F (3,22) = 1.267 | 0.449 | ns |
| | | P18-20 | -24.4 ± 2.91 | 9.65 | -31.6 ± 3.77 | 12.5 | | | 0.346 | ns |
| | Slope | P12-13 | 29.4 ± 0.68 | 1.67 | 25.8 ± 0.34 | 0.96 | | F (3,22) = 5.996 | 0.200 | ns |
| | | P18-20 | 33.3 ± 1.85 | 6.14 | 27.3 ± 1.02 | 3.37 | | | 0.004 | ** |
| Palm3 Hets | OHC length | OHC1 | 15.1 ± 1.05 | 2.56 | 12.8 ± 0.99 | 2.41 | Two-way ANOVA with Holm-Šídák's multiple comparisons correction | F (1,32) = 16.46 | 0.0009 | *** |
| | | OHC2 | 15.1 ± 1.19 | 2.92 | 11.7 ± 0.73 | 1.79 | | | 0.0009 | *** |
| | | OHC3 | 16.2 ± 1.14 | 2.79 | 11.9 ± 0.98 | 2.41 | | | 0.0009 | *** |

**Supplementary Table 2: Statistics summary II**

| | | | Contralateral | | Ipsilateral | | Statistical test | F value | *P* value | Significance |
|---|---|---|---|---|---|---|---|---|---|---|
| | | | Mean ± SEM | SD | Mean ± SEM | SD | | | | |
| Rescue | ABR wave I-V amplitude | 60 | 0.30 ± 0.00 | 0.00 | 0.93 ± 0.32 | 1.02 | Two-way ANOVA with Holm-Šídák's multiple comparisons correction | F( (1,94) = 16.12 | 0.0006 | *** |
| | | 70 | 0.72 ± 0.42 | 1.34 | 1.93 ± 0.71 | 2.25 | | | 0.0006 | *** |
| | | 80 | 1.56 ± 0.61 | 1.92 | 3.37 ± 0.88 | 2.78 | | | 0.0006 | *** |
| | | 90 | 2.73 ± 0.95 | 3.00 | 5.41 ± 0.76 | 2.41 | | | 0.0006 | *** |
| | | 100 | 5.98 ± 0.72 | 2.29 | 8.23 ± 0.82 | 2.59 | | | 0.0006 | *** |
| | ABR threshold | 6 | 82.0 ± 2.49 | 7.89 | 75.0 ± 2.24 | 7.07 | Two-way ANOVA with Holm-Šídák's multiple comparisons correction | F (1,75) = 13.24 | 0.0020 | ** |
| | | 12 | 79.0 ± 2.77 | 8.76 | 74.0 ± 2.21 | 6.99 | | | 0.0020 | ** |
| | | 24 | 88.0 ± 1.33 | 4.22 | 82.0 ± 2.00 | 6.33 | | | 0.0020 | ** |
| | | click | 87.0 ± 3.67 | 11.6 | 77.0 ± 4.23 | 13.4 | | | 0.0020 | ** |
| | DPOAE intensity | 5.67 kHz | -15.7 ± 0.88 | 2.79 | -15.5 ± 0.68 | 2.14 | Two-way ANOVA Mixed-effects analysis with Holm-Šídák's multiple comparisons correction | not significant | | |
| | | 11.33 kHz | -14.5 ± 2.29 | 7.26 | -12.7 ± 0.2.75 | 8.69 | | | | |
| | | 22.67 kHz | -18.8 ± 2.15 | 6.79 | -18.7 ± 0.59 | 1.87 | | | | |
| | Transduction rate | IHC | 9.29 ± 2.65 | 7.96 | 80.94 ± 7.87 | 23.61 | Two-way ANOVA with Holm-Šídák's multiple comparisons correction | F (1,32) = 220.4 | <0.0001 | **** |
| | | OHC row 1 | 3.55 ± 1.41 | 4.22 | 68.84 ± 8.62 | 25.86 | | | <0.0001 | **** |
| | | OHC row 2 | 3.35 ± 1.61 | 4.84 | 60.42 ± 9.77 | 29.32 | | | <0.0001 | **** |
| | | OHC row 3 | 0.00 ± 0.00 | 0.00 | 63.33 ± 7.99 | 23.98 | | | <0.0001 | **** |
| | Hair cell survival | OHC row 1 | 19.56 ± 2.28 | 6.84 | 29.89 ± 3.04 | 9.13 | Two-way ANOVA with Holm-Šídák's multiple comparisons correction | F (1,24) = 17.47 | 0.001 | ** |
| | | OHC row 2 | 15.11 ± 1.95 | 5.84 | 21.33 ± 2.46 | 7.37 | | | 0.040 | * |
| | | OHC row 3 | 23.44 ± 1.80 | 5.41 | 25.00 ± 1.54 | 4.61 | | | 0.539 | ns |
| | Length | OHC row 1 | 7.77 ± 0.25 | 0.74 | 13.8 ± 0.58 | 1.74 | Two-way ANOVA with Holm-Šídák's multiple comparisons correction | F (1,24) = 277.1 | <0.0001 | **** |
| | | OHC row 2 | 7.84 ± 0.38 | 1.14 | 14.1 ± 0.39 | 1.18 | | | <0.0001 | **** |
| | | OHC row 3 | 7.20 ± 0.23 | 0.69 | 14.4 ± 0.53 | 1.59 | | | <0.0001 | **** |

#### Supplementary Table 3: Statistics summary III

| | | | Median (IQR) | | Statistical test | *P* value | Significance |
|---|---|---|---|---|---|---|---|
| | | | WT | *Palm3* -KO | | | |
| Tomo-SSC | no. SSC/membrane | | 2.72 (1.84 - 3.37) | 0.00 (0.00 - 0.00) | Two-tailed Mann-Whitney test | <0.0001 | **** |
| | avg. distance SSC-PM | | 34.0 (27.2 - 43.3) | 56.6 (25.4 - 82.7) | | 0.0015 | ** |
| | surface/volume mesh | | 0.33 (0.32 - 0.44) | 0.20 (0.12 - 0.24) | | 0.0056 | ** |
| | avg. distance mito-PM | | 292.6 (122.5 - 1004) | 831.2 (358.3 - 1383) | | <0.0001 | **** |
| αII-Spectrin | norm. avg. intensity | OHC lateral wall | 0.99 (0.92 - 1.08) | 0.50 (0.42 - 0.62) | Two-tailed Mann-Whitney test | <0.0001 | **** |
| | | OHC cuticular plate | 1.00 (0.98 - 1.03) | 1.04 (0.87 - 1.67) | | 0.0075 | ** |
| βI-Spectrin | norm. avg. intensity | OHC lateral wall | 0.99 (0.95 - 1.05) | 0.18 (0.12 - 0.20) | | <0.0001 | **** |
| | | OHC cuticular plate | 0.99 (0.93 - 1.08) | 1.11 (0.96 - 1.28) | | <0.0001 | **** |
| βII-Spectrin | norm. avg. intensity | OHC lateral wall | 0.99 (0.91 - 1.07) | 0.81 (0.67 - 0.98) | | <0.0001 | **** |
| | | OHC cuticular plate | 0.99 (0.93 - 1.05) | 0.75 (0.59 - 1.06) | | <0.0001 | **** |
| Palm3 Hets | Palm3 norm. avg. intensity | | 1.02 (0.96 - 1.07) | 0.77 (0.65 - 1.04) | Two-tailed Mann-Whitney test | <0.0001 | **** |
| Rescue | | | **non-transduced** | **Palm3-transduced** | | | |
| | αII-Spectrin | norm. avg. intensity | 1.02 (0.89 - 1.10) | 1.62 (1.34 - 2.34) | Two-tailed Mann-Whitney test | <0.0001 | **** |
| | Prestin | norm. avg. intensity | 1.01 (0.97 - 1.03) | 0.77 (0.71 - 0.87) | | <0.0001 | **** |
| | | coefficient of variation | 0.46 (0.38 - 0.56) | 0.28 (0.25 - 0.32) | | 0.0002 | *** |
| | | | **ipsilateral** | **contralateral** | | | |
| | IHC count | | 37.0 (34.5 - 38.5) | 35.0 (27.5 - 36.0) | Two-tailed Wilcoxon matched-pairs signed rank test | 0.3502 | ns |
